# Mapping the Transcriptional Landscape of Drug Responses in Primary Human Cells Using High-Throughput DRUG-seq

**DOI:** 10.1101/2025.06.03.657593

**Authors:** Lauren Baugh, Sébastien Vigneau, Srijani Sridhar, Sarah Boswell, George Pilitsis, John Bradley, Olga Allen, Laura Rand, Amanda Riccardi, Kelsey Roach, Kevin Lema, Ayla Ergun

## Abstract

To advance our understanding of drug action in physiologically-relevant systems, we developed a high-throughput transcriptomic atlas of compound responses in primary human cell types. Leveraging the scalable and cost-effective Digital RNA with the pertUrbation of Genes (DRUG-seq) assay, we profiled gene expression responses to 89 pharmacologically-active compounds across six concentrations in four distinct primary cell types: aortic smooth muscle cells (AoSMCs), skeletal muscle myoblasts (SkMMs), dermal fibroblasts, and melanocytes. Through rigorous quality control and normalization, we generated reproducible and cell type-resolved transcriptomic signatures, enabling the discovery of both shared and divergent regulatory programs.

This dataset revealed core cellular responses, such as brefeldin A-mediated ER stress across all cell types, as well as lineage-specific effects, including dexamethasone-induced hypoxia signaling in AoSMCs, complex inflammatory responses linked to epithelial-to-mesenchymal transition pathways in SkMMs, TGF-β-modulated states in fibroblasts, and dabrafenib-driven transcriptional shifts towards quiescence in melanocytes. By integrating systematic perturbations with primary models, this dataset serves as a resource for building systems-level models of drug response and mechanism. Ultimately, we aim to accelerate predictive pharmacology by enabling high-throughput data generation grounded in human biology and readily usable by artificial intelligence models.

## 1. Introduction

Combining high-throughput screening (HTS) with transcriptomic profiling has become a powerful approach in drug discovery, enabling cheaper and faster genome-wide measurement of cellular responses to perturbations. Multiple transcriptional profiling assays such as RASL-seq, TempO-seq, Plate-seq, and Digital RNA with pertUrbation of Genes (DRUG-seq) have been developed in order to address this need (Li et al., 2012; Bundy et al., 2024; Bush et al., 2017; Ye et al., 2018). Compared to traditional bulk RNA-seq, DRUG-seq uses smaller reaction volumes and shallow sequencing, enabling smaller well formats (such as 384-well plates) and more cost-effective high-throughput profiling. The scalability and precision of DRUG-seq make it a powerful strategy for both early-stage drug discovery and the functional characterization of chemical perturbations in diverse cellular contexts. This approach is particularly valuable for mechanisms of action (MOA) studies, as it enables the correlation of many perturbations with cellular transcriptomic changes, providing insights into affected signaling pathways and further aiding the identification of drug classes with shared mechanisms (Lamb et al., 2006; Subramanian et al., 2017).

In addition to the advantages of performing HTS coupled with DRUG-seq, the choice of disease-relevant models is also critical in such perturbation studies. While immortalized cell lines are widely used in high-throughput drug screening due to their ease of culture and scalability, they often fail to accurately represent *in vivo* human biology (Voloshin et al., 2023). Primary human cells, on the other hand, retain the genetic, epigenetic, and functional characteristics of their tissue of origin, making them a more physiologically-relevant model for studying drug responses (Dai et al., 2024). These characteristics are highly relevant for drug discovery, as compounds that appear effective in cell lines often fail in pre-clinical studies due to unrealistic responses or missing key signaling pathways (Eastman, 2016). By leveraging primary human cells in HTS, more representative transcriptomic and phenotypic data can be generated, contributing to a better understanding of drug mechanisms, the development of more accurate predictive models, and fewer late-stage failures in drug development.

We present a first-of-its-kind, high-throughput DRUG-seq study in four human primary cell types: aortic smooth muscle cells (AoSMCs), skeletal muscle myoblasts (SkMMs), dermal fibroblasts, and melanocytes. The cell types were chosen to provide a diverse set of cell types and phenotypes and demonstrate that DRUG-seq is a powerful HTS tool for use with various disease-relevant tissues. In each of these cell types, we tested 89 pharmacologically-active compounds at six concentrations (9.5, 28.5, 95, 300, 900, 3000 nM) with four replicates and DRUG-seq readout, resulting in 11,808 transcriptomes. To reduce cost and achieve high-throughput, experiments were conducted in a 384-well plate format using a scalable automated workflow (Figure 1). In this study, we demonstrate that our DRUG-seq workflow captures transcriptomic profiles in primary cells with sufficient resolution to characterize dose-dependent gene-and pathway-level responses to a large number of compounds.

**Figure 1:**
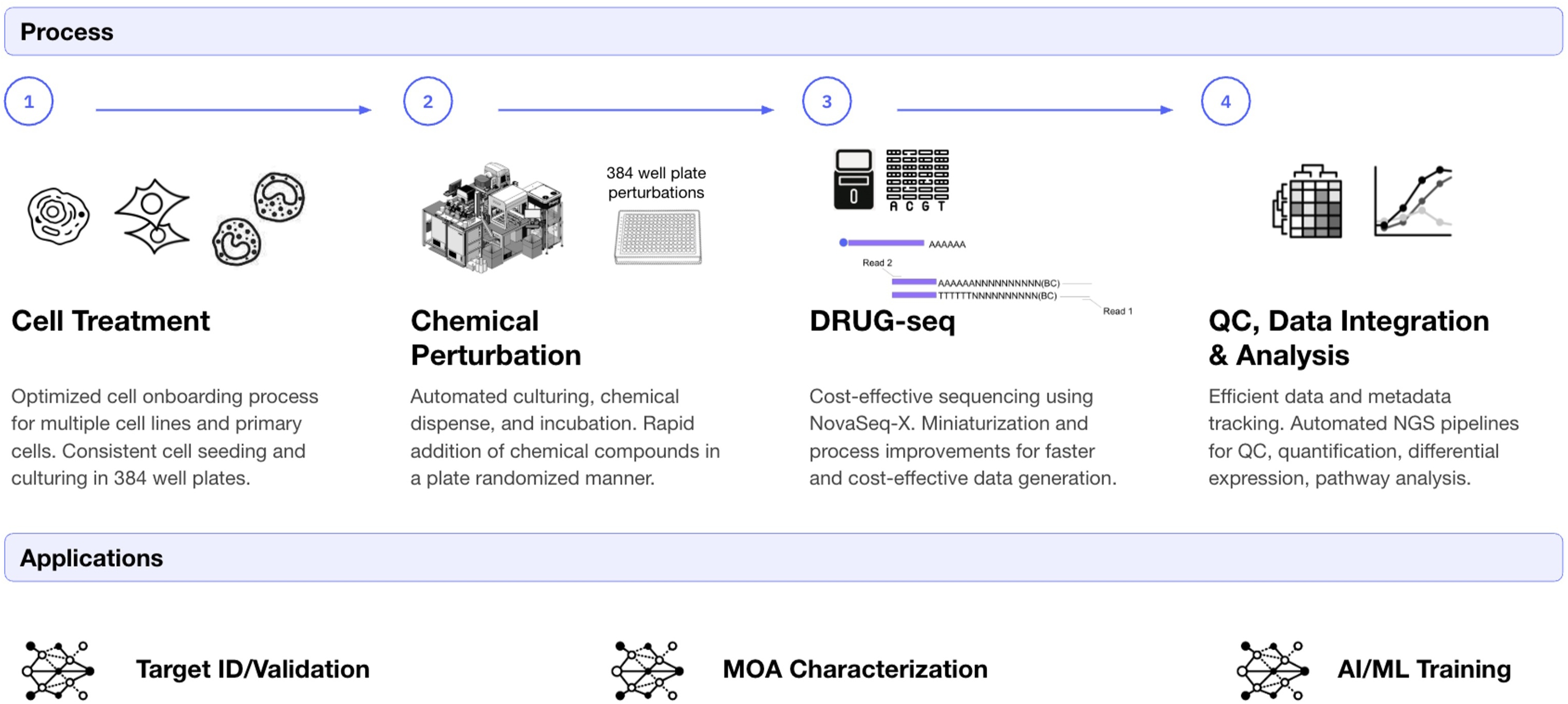
Arrayed DRUG-seq chemical perturbation workflow.

## 2. Results

### 2.1 Experimental design

Human primary AoSMCs, SkMMs, dermal fibroblasts, and melanocytes were subjected to 24 hours of chemical treatment using 89 selected compounds at six different concentrations, as described in Table 1. Compounds were selected from an in-house small molecule library comprising both curated and commercially available pharmacologically active agents, chosen to span a broad range of known MOAs. Each plate included three positive controls known to elicit transcriptional changes—trichostatin A (TSA), brefeldin A (BFA), and dexamethasone (DEX)—and a vehicle control, dimethyl sulfoxide (DMSO). The positive controls were selected based on their known broad transcriptomic effects. TSA is a histone deacetylase (HDAC) inhibitor, leading to increased histone acetylation and transcription (Yoshida et al., 1990). Similarly broad, BFA causes the collapse of the Golgi apparatus, which prevents newly synthesized proteins from being transported; vesicles and proteins begin to accumulate and overwhelm the ER folding capacity, which further causes a cascading effect on multiple pathways (Fujiwara et al., 1988). DEX is a synthetic glucocorticoid that mimics the effects of cortisol, a hormone produced by the adrenal glands. It binds to the glucocorticoid receptor (GR) and regulates gene expression, leading to widespread effects on inflammation, metabolism, and immune responses (Oakley & Cidlowski, 2013; Rhen & Cidlowski, 2005). These controls were used as a baseline for our QC metrics.

**Table 1:** 86 pharmacologically-active compounds plus controls were tested at 6 concentrations (9.5, 28.5, 95, 300, 900, 3000 nM) using four technical replicates in 4 human primary cell types.

| Treatment Name | Replicates per Dose | Concentration Range | Function |
| --- | --- | --- | --- |
| DMSO | 6 | 0.0625%, 0.1875%, 0.625% | Vehicle control, added at different volumes to match drug concentrations. |
| brefeldin A | 4 | 9.5, 28.5, 95, 300, 900,3000 nM | Positive control of transcriptional response.<br>ER-Golgi trafficking inhibitor |
| dexamethasone | 4 | 9.5, 28.5, 95, 300, 900,3000 nM | Positive control of transcriptional response.<br>Anti-inflammatory |
| trichostatin A | 4 | 9.5, 28.5, 95, 300, 900,3000 nM | Positive control of transcriptional response.<br>HDAC inhibitor |
| 86 Test compounds | 4 | 9.5, 28.5, 95, 300, 900,3000 nM | 11 compounds per plate<br>(8 compounds on last plate) |

Following sequencing and alignment, QC metrics were evaluated. In particular, samples were sequenced to a depth of at least 2 million reads, and all controls for all cell types had an average recovery of just over 10,000 gene transcripts (including noncoding and protein-coding genes and pseudogenes) with greater than three unique UMI counts (Supplementary Figure 1). Additionally, all samples for each cell type showed a tight distribution across all QC metrics.

To assess experimental consistency and reproducibility across experimental conditions, we compared transcriptional profiles between technical replicates and across different cell types. Gene expression profiles across plates revealed high plate-to-plate consistency within each cell type for all controls, including transcriptomic response to DMSO and the positive controls (TSA, BFA, and DEX). In particular, control replicates across plates of the same cell type consistently showed a median Pearson correlation of gene expression profiles greater than 0.99 (Figure 2*a*), whereas comparisons between plates containing different cell types showed markedly lower correlations with a median value of less than 0.79. These findings confirmed that transcriptional profiles are highly reproducible within individual cell types while maintaining a clear separation between distinct cell populations.

**Figure 2:**
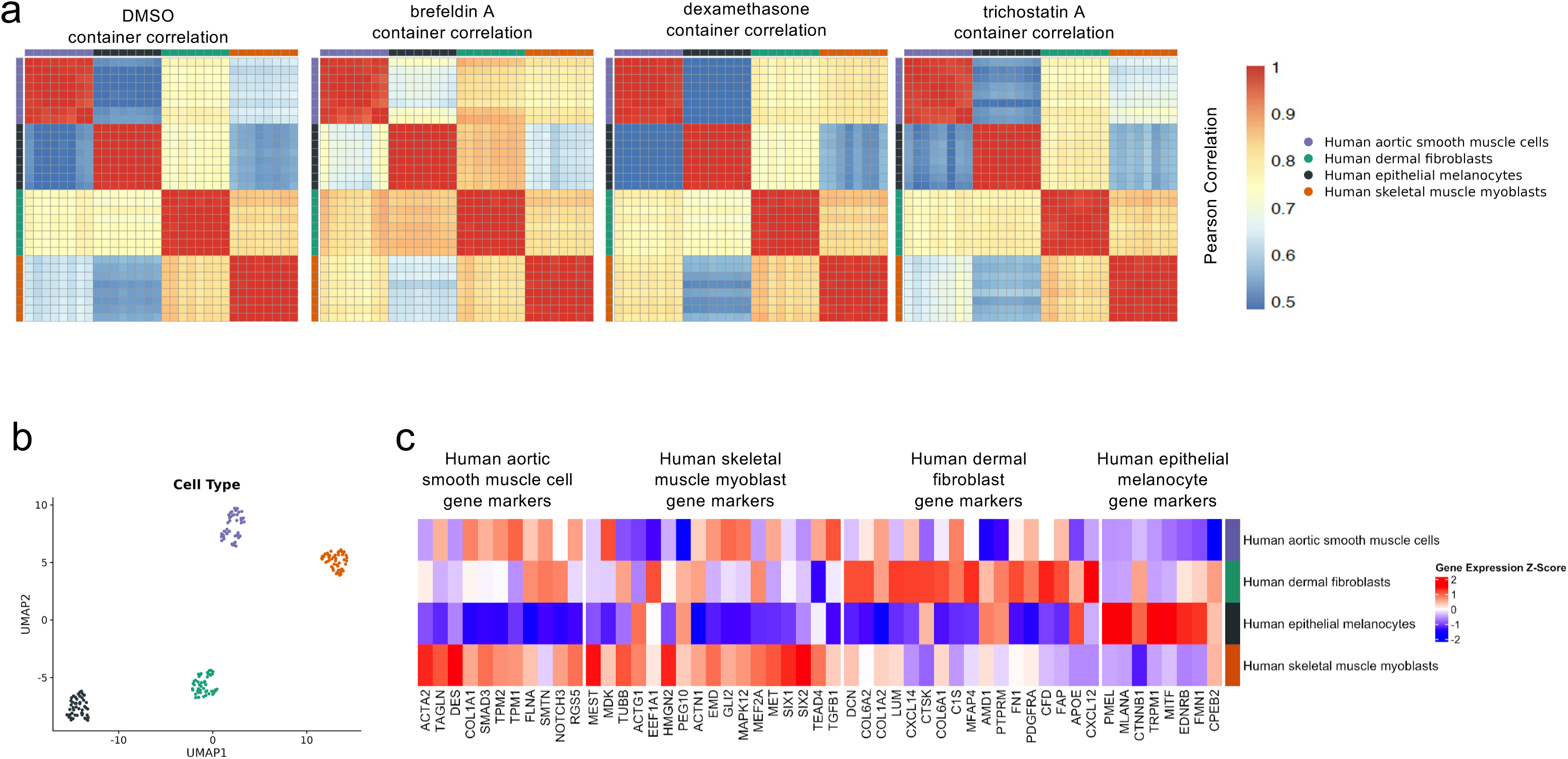
Plate-to-plate correlation across all cell types for known cell type specific gene markers. *(a)* The average genome-wide Pearson’s correlation value was calculated across the wells containing either DMSO or the positive controls wells (brefeldin A, dexamethasone, and trichostatin A) for each pair of plates. Plates with the same cell type correlate strongly, with a median coefficient value >0.99, while plates with different cell types show weaker correlation. *(b)* A UMAP projection of gene expression data for wells containing only the lowest DMSO concentration for each cell type. *(c)* Mean gene expression data for a curated list of cell type-specific gene markers, z-score normalized for each marker gene across cell types.

### 2.2 DRUG-seq data capture known cell type differences

To confirm that DRUG-seq can unambiguously distinguish between cell types, wells treated with the lowest (0.0625%) DMSO concentration were compared. Uniform Manifold Approximation and Projection (UMAP) of the lowest-concentration DMSO controls distinctly separated AoSMCs, SkMMs, dermal fibroblasts, and melanocytes into four non-overlapping clusters (Figure 2*b*).

Next, we mapped literature-curated cell type markers, as described in the methods, against our DMSO-treated primary cell types (Figure 2*c*). As expected, dermal fibroblasts and melanocytes exhibited distinct marker gene expression from each other and from AoSMCs and SkMMs, whereas AoSMCs and SkMMs expression profiles were more similar to each other, consistent with their lineage and phenotypic proximity. Fibroblasts exhibited strong expression of extracellular matrix (ECM) related genes such as *COL1A2*, *COL6A1/2*, *DCN*, and *FAP*, alongside stromal regulators like *PDGFRA*. Melanocytes, on the other hand, expressed pigment synthesis genes (*PMEL* and *MLANA*) and lineage-determining factors such as *MITF*. AoSMCs and SkMMs shared expression of genes encoding contractile machinery, including *TAGLN*, consistent with their roles in tension generation, but differed in the expression of other lineage-specific markers. For instance, AoSMCs were enriched in smooth muscle-specific and differentiation genes such as *SMTN* and *TPM1*, as well as ECM regulators like *COL1A1*. Similarly, SkMMs uniquely expressed the myogenic proliferation-associated gene *PEG10*.

Together, these findings highlighted DRUG-seq’s power to resolve both transcriptome-wide and gene-level differences between human primary cell types, demonstrating clear marker specificity for melanocytes and dermal fibroblasts, while capturing the more subtle, transcriptional distinctions between AoSMCs and SkMMs.

### 2.3 Effect of pharmacological perturbations on primary cell types

In order to reveal the transcriptional response of primary human AoSMCs, SkMMs, dermal fibroblasts, and melanocytes to chemical perturbations, we performed an unbiased analysis after normalizing the data for potential confounding factors (Methods). First, we performed graph-based clustering and UMAP visualization of samples based on their transcriptional profiles. Second, in order to gain insight into the induced pathway changes, we performed gene set enrichment analysis (GSEA) on the MSigDB Hallmark gene sets (Liberzon et al., 2015). Third, we identified dose-dependent gene-expression changes for every compound and cell type (Methods).

#### Shared compound responses across cell types

A global analysis confirmed that our positive-control compounds (TSA, BFA, and DEX) induced strong, dose-dependent transcriptomic changes, as seen in PCA plots where the first two components accounted for 33–42% of total variance (Supplementary Figure 2). TSA and BFA clustered separately across all cell types, reflecting their distinct mechanisms: chromatin remodeling by TSA and Golgi apparatus disruption by BFA. TSA showed clear dose-dependent divergence from DMSO, consistent with its broad HDAC inhibition (Figures 3–6, panel *a*). BFA caused consistent, widespread perturbations across all four cell types, particularly at mid-to-high doses, including downregulation of ER–Golgi transport and protein synthesis pathways, based on the MSigDB Hallmark gene sets (Figures 3–6, panels *a* and *b*; Fujiwara et al., 1988). In contrast, DEX triggered transcriptional changes specifically in AoSMCs, with little effect in other cell types (Supplementary Data), indicating a cell type-specific response likely mediated by glucocorticoid receptor signaling. Overall, higher compound concentrations amplified the inter-cluster distances, demonstrating dose-dependent transcriptional perturbation.

**Figure 3:**
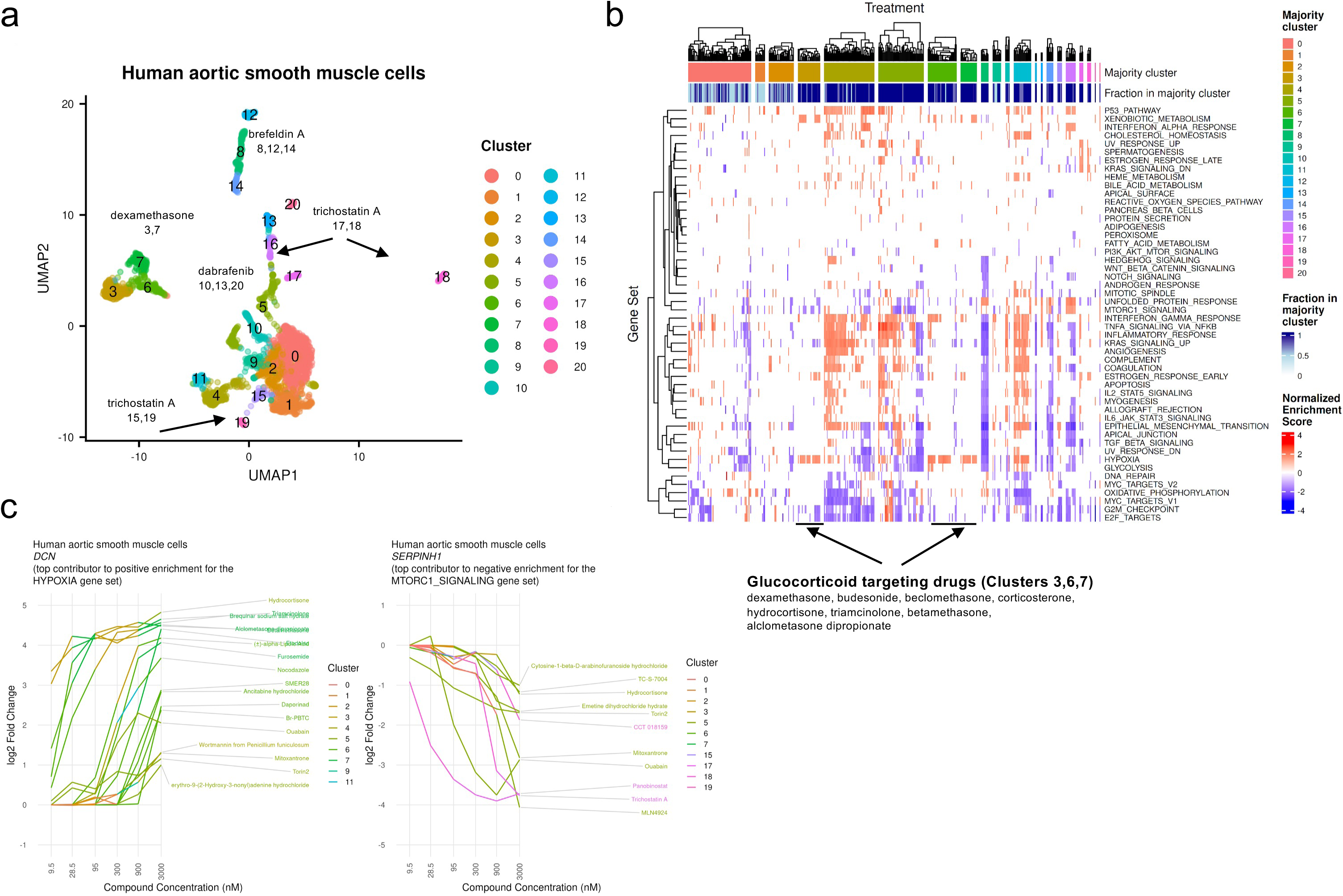
Gene set enrichment analysis and UMAP representation for human aortic smooth muscle cells. *(a)* UMAP projection of all compounds and all concentrations with control clusters labeled (brefeldin A, dexamethasone, and trichostatin A) and dabrafenib. *(b)* MsigDB hallmark gene set enrichment analysis for clusters displayed in the UMAP projection, highlighting a subset of clusters and compounds of interest. *(c)* Example genes in enriched pathways with a dose-dependent response.

**Figure 4:**
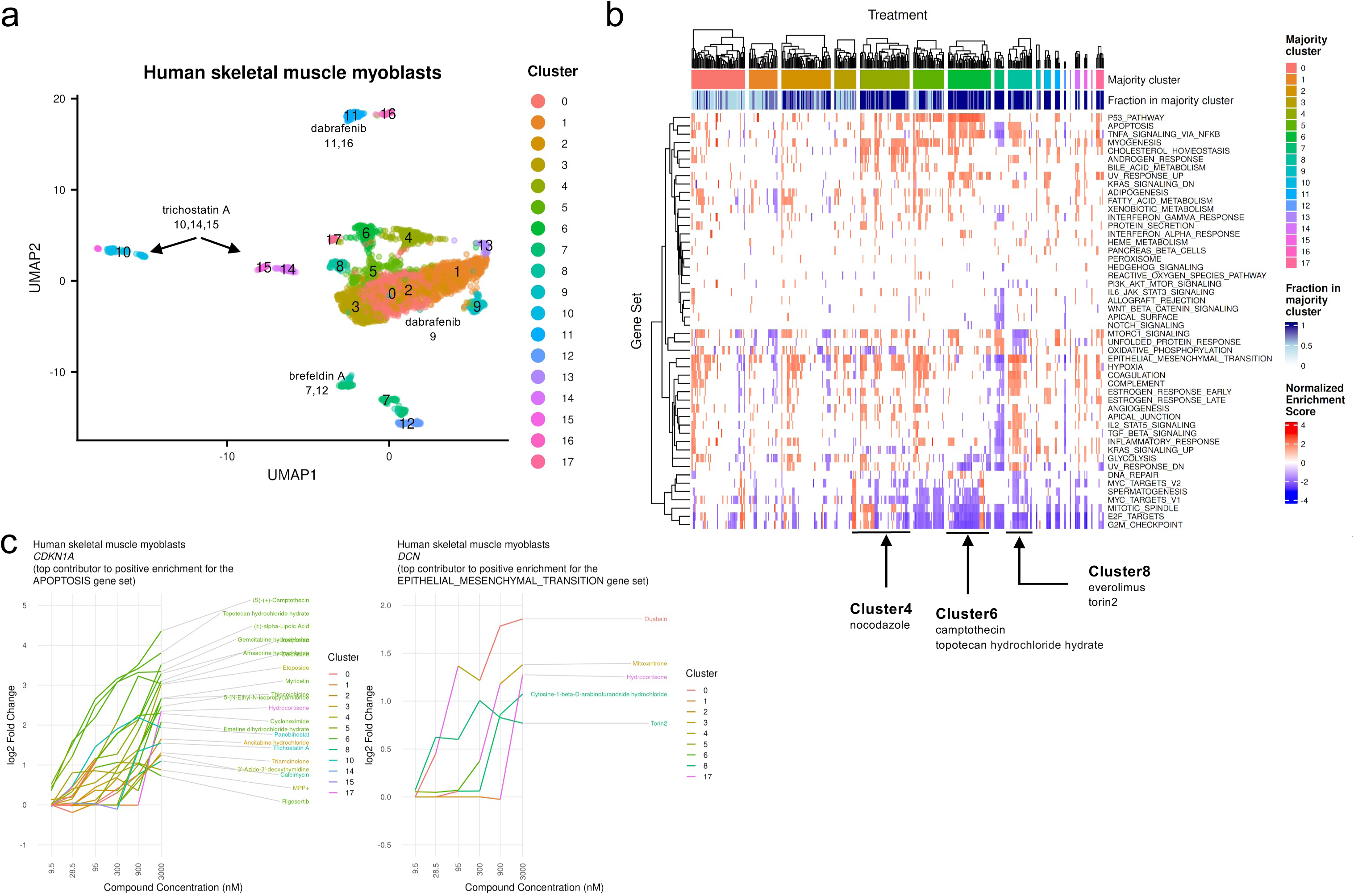
Gene set enrichment analysis and UMAP representation for human skeletal muscle myoblasts. *(a)* UMAP projection of all compounds and all concentrations with control clusters labeled (brefeldin A, dexamethasone, and trichostatin A) and dabrafenib. *(b)* MsigDB hallmark gene set enrichment analysis for clusters displayed in the UMAP projection, highlighting a subset of clusters and compounds of interest. *(c)* Example genes in enriched pathways with a dose-dependent response.

**Figure 5:**
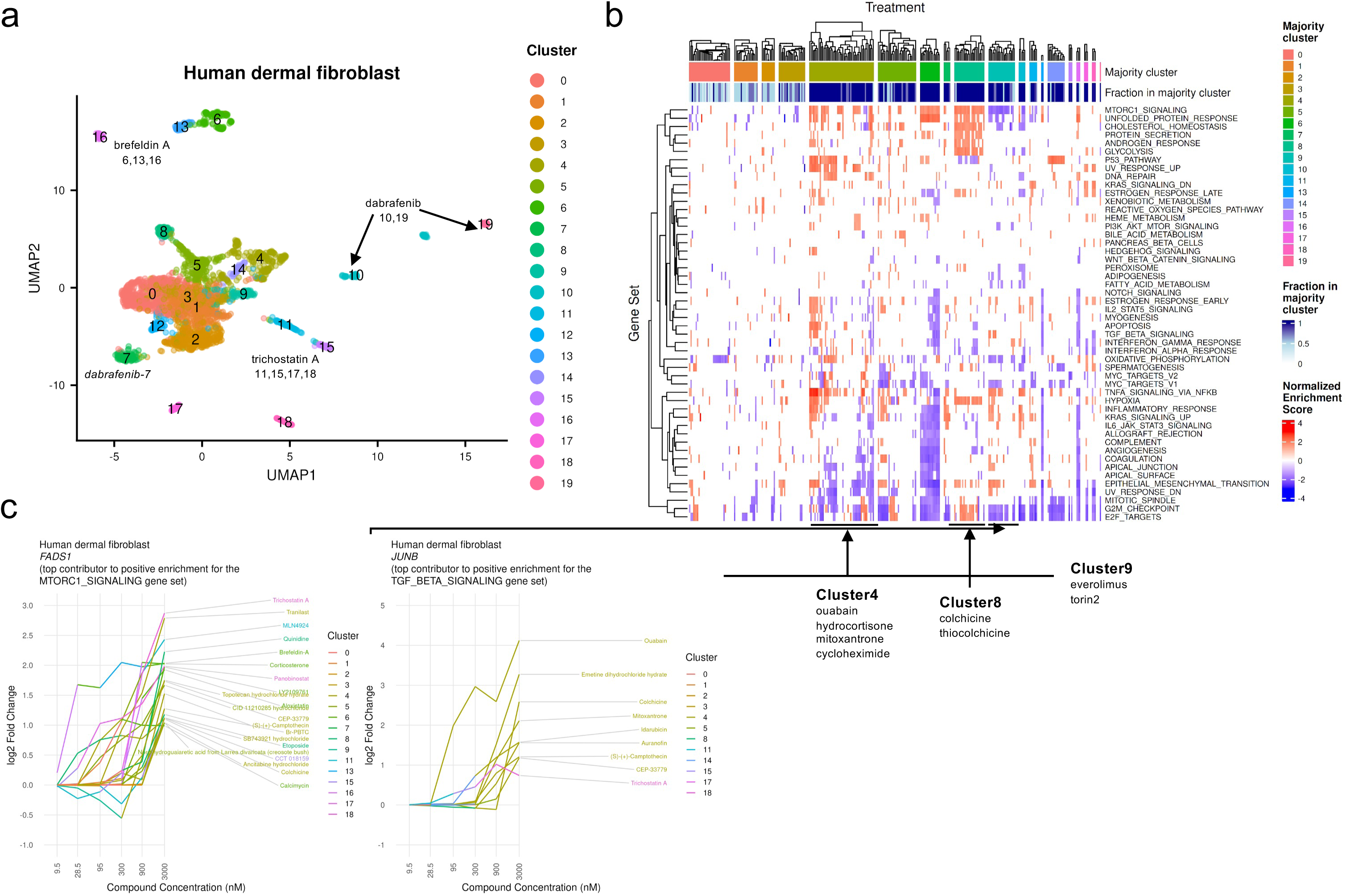
UMAP and gene set enrichment analysis for human dermal fibroblasts. *(a)* UMAP projection of all compounds and all concentrations clustered with control clusters labeled (brefeldin A, dexamethasone, and trichostatin A) and dabrafenib. *(b)* MsigDB hallmark gene set enrichment analysis for clusters displayed in the UMAP projection, highlighting a subset of clusters and compounds of interest. *(c)* Example genes in enriched pathways with a dose-dependent response.

**Figure 6:**
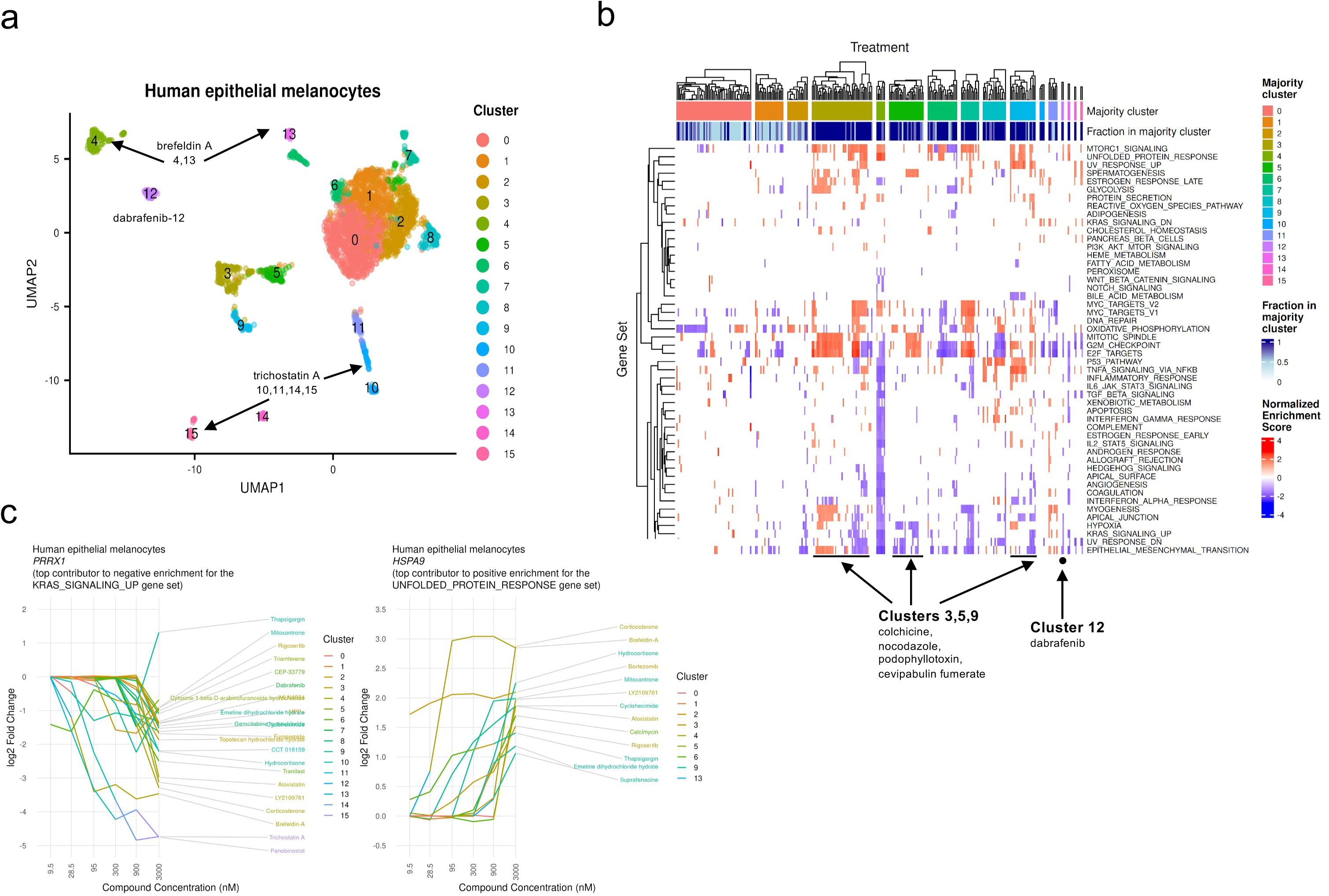
UMAP and gene set enrichment analysis for human epithelial melanocytes. *(a)* UMAP projection of all compounds and all concentrations clustered with control clusters labeled (brefeldin A, dexamethasone, and trichostatin A) and dabrafenib. *(b)* MsigDB hallmark gene set enrichment analysis for clusters displayed in the UMAP projection, highlighting a subset of clusters and compounds of interest. *(c)* Example genes in enriched pathways with a dose-dependent response.

Outside of the control clusters, dabrafenib was the only compound to elicit consistently strong transcriptional responses across all four primary human cell types. Although primarily developed to inhibit mutant *BRAF* V600E in cancers, dabrafenib has been shown to influence basal MAPK pathway activity even in *BRAF* wild-type cells, leading to downstream effects on gene expression, proliferation, and stress responses (Bowyer et al., 2015). In Figures 3–6 (panel *b*), clusters containing dabrafenib consistently impacted KRAS signaling and epithelial-to-mesenchymal transition (EMT) pathways, aligning with the established role of MAPK/ERK signaling in maintaining mesenchymal and migratory phenotypes (King et al., 2013).

#### Dexamethasone induces cell type-specific transcriptional responses in AoSMCs

In AoSMCs, DRUG-seq profiling of our compound library revealed multiple distinct phenotypes (Figure 3*a*). As mentioned above, we observed a strong response to BFA in all cell types, including AoSMCs, as indicated by the broad downregulation of gene sets involved in inflammatory pathways, cell differentiation, and morphogenesis (clusters 8, 12, and 14, Figure 3*a,b*). Interestingly, DEX had a cell type-specific effect on AoSMCs. DEX treatment at multiple concentrations induced similar genome-wide profiles and converged in clusters 3 and 7, alongside other compounds known to be corticosteroids or GR agonists. Additionally, cluster 6 included several other corticosteroids or GR agonists, which all together lead to similar pathway enrichment profiles (Figure 3*a,b*). These compound clusters lead to a transcriptional state in AoSMCs characterized by upregulation of hypoxia-responsive genes and suppression of mTORC1 signaling. This is consistent with the well-documented ability of glucocorticoids to induce catabolic and stress-adaptive programs, reduce protein synthesis, and modulate oxygen-sensing pathways (Reil et al., 1999). In other cell types, DEX does not lead to differential expression changes and has a similar transcriptomic profile to DMSO (Supplementary Table 8).

To further confirm the effects of the corticosteroids and GR agonist compounds in a dose-dependent manner, hypoxia pathway activation is represented by upregulation of *DCN* (Figure 3*c*), a gene involved in ECM organization and known to be induced under hypoxic and stress conditions. Conversely, *SERPINH1*, which encodes HSP47, an ER-resident collagen chaperone essential for collagen folding and secretion, was downregulated in a dose-dependent manner. Given that HSP47 expression supports the high biosynthetic and secretory demands of anabolic cells, its suppression is consistent with a reduction in protein synthesis capacity and secretory pathway activity. This downregulation aligns with the attenuation of mTORC1 signaling observed in these clusters, as mTORC1 is a key driver of translational output and ER folding demand, and its inhibition would be expected to reduce the need for chaperone-mediated protein quality control.

#### Epithelial-mesenchymal transition pathway is activated in SkMMs

For SkMMs DRUG-seq data, UMAP-resolved CCA clusters mapped to distinct cellular differentiation pathways. BFA mirrored its effects in AoSMCs. Clusters 7 and 12 (Figure 4*a,b*) showed downregulation of protein transport genes and ER stress markers, indicative of disrupted secretory pathways essential for muscle regeneration. Clusters 4, 6, and 8 (Figure 4*b*) highlighted interesting biology around the known phenotype plasticity and mesenchymal characteristics also found in myoblasts (Cho & Doles, 2017). While all three of these clusters were downregulated for pathways involving cell cycling and enriched for several inflammatory pathways, clusters 4 and 8 were also enriched for the EMT pathway. The enrichment of the EMT pathway in clusters 4 and 8 was likely caused by the undifferentiated, transcriptionally plastic state of the SkMMs. When exposed to stressors such as nocodazole, which has been shown to promote EMT in epithelial cells (cluster 4), or mTOR/epigenetic inhibitors like everolimus and torin2 (cluster 8), these cells diverted away from myogenesis and instead engaged in EMT-like transcriptional programs characteristic of repair, fibrosis, or developmental arrest (Kawami et al., 2019).

Interestingly, cluster 6 compounds arrested the cell cycle and suppressed anabolic processes similarly to cluster 4, but without triggering transcriptional plasticity or mesenchymal reprogramming. Unlike clusters 4 and 8, which featured inflammatory environments, cluster 6 compounds, which included known cell cycle arrest compounds camptothecin and topotecan, enforced a cytostatic state. This state reflected a different cellular decision point in undifferentiated SkMMs: one focused on damage containment rather than fate change (Czuwara-Ladykowska et al., 2001; Feeney et al., 2003). Given that successful cancer therapeutics look to avoid inducing EMT, screening techniques, like ours, that capture EMT pathway signatures can be very valuable (Lamouille et al., 2014). SkMMs are uniquely positioned for this type of screening since they are distinct from fully epithelial or mesenchymal lines and can modulate EMT without fully inducing it (Dongre & Weinberg, 2019). By capturing high resolution transcriptomic signatures with DRUG-seq, our data pointed to compounds that capture these transitional cell states and differentiation pathways.

Figure 4*c* highlights representative genes with clear dose–response relationships that reinforce pathway enrichments observed in clusters 4, 6, and 8. The upregulation of *DCN*, an extracellular matrix proteoglycan known to modulate TGF-β signaling and promote mesenchymal traits, is consistent with EMT pathway enrichment in clusters 4 and 8. In contrast, the strong dose–responsive increase in *CDKN1A*, a cyclin-dependent kinase inhibitor that leads to cell cycle arrest and is a key mediator of apoptosis, reinforced the cell cycle arrest and death-associated phenotype of cluster 6.

#### TGF-β signaling is induced in dermal fibroblasts

DRUG-seq revealed pronounced sensitivity of dermal fibroblasts to BFA, as evidenced by the clusters 6, 13, and 16 (Figure 5*a*), demonstrating robust upregulation of protein secretion pathways (Figure 5*b*). The universal impact of BFA across cell types highlighted its utility as a broad secretory pathway inhibitor. DRUG-seq also captured different fibroblast phenotypes in response to different categories of compounds. Cluster 8 exhibited coordinated upregulation of hypoxia, glycolysis, protein secretion, unfolded protein response (UPR), and mTORC1 signaling pathways, indicating a metabolically-reprogrammed phenotype characteristic of fibrotic processes. Colchicine and thiocolchicine (a derivative of colchicine) clustered together, and colchicine has been shown to induce an ECM-producing myofibroblast phenotype, further validating this finding (Sandbo et al., 2013). The concurrent activation of these pathways reflects increased protein synthesis and secretory demand consistent with matrix-producing myofibroblasts, as fibrogenic signals trigger metabolic rewiring to meet the high bioenergetic demands of dermal fibroblasts.

Conversely, compounds in cluster 9, including mTOR inhibitors everolimus and torin2, suppressed mTORC1 signaling and UPR while upregulating TGF-β signaling. This observation aligns with research demonstrating that mTOR inhibition affects glycolytic enzymes and mitochondrial function in fibroblasts (Shin et al., 2025). TGF-β signaling was also upregulated in cluster 4, specifically in response to different concentrations of ouabain, which has previously been shown to influence TGF-β response (La et al., 2016). Other compounds in cluster 4 included GR agonists and antagonists, which produced a cell type-specific effect in fibroblasts. Unlike in AoSMCs, where GR agonists such as dexamethasone induced a catabolic, stress-adaptive program, GR antagonist treatment in fibroblasts led to the upregulation of mTORC1 and hypoxia-related pathways, a pro-anabolic, potentially fibrogenic state. This divergence suggests that GR signaling plays distinct regulatory roles depending on cellular context, potentially due to baseline differences in chromatin accessibility, GR cofactor availability, or crosstalk with TGF-β signaling pathways (Butz et al., 2023).

Figure 5*c* provides a quantitative view of pathway-specific gene expression changes across dose gradients in dermal fibroblasts. Upregulation of *FADS1*, a lipid biosynthesis enzyme downstream of mTORC1, suggests a matrix-producing myofibroblast phenotype characterized by anabolic and secretory activity. In contrast, upregulation of *JUNB*, a component of the AP-1 transcription factor complex and a known effector of TGF-β signaling, indicates activation of early transcriptional programs involved in fibrotic remodeling and stress response.

#### BRAF-inhibitor dabrafenib shows expected mechanisms of action in melanocytes

In the melanocyte DRUG-seq transcriptomic data, BFA showed similar patterns as in the previous cell types (Figure 6*a*, clusters 4 and 13). Additionally, dabrafenib-treated samples across concentrations were marked by robust downregulation of myogenesis, apical junctions, EMT, and KRAS signaling, which are hallmarks of suppressed differentiation and epithelial identity (cluster 12). This was coupled with a lack of compensatory upregulation in proliferative programs, suggesting that dabrafenib-induced BRAF inhibition in non-transformed melanocytes lead to a checkpoint-engaged, quiescent phenotype rather than hyperproliferation, as seen previously in melanoma cells treated with dabrafenib (Yang et al., 2021). This mirrors known BRAF/MAPK pathway suppression effects in melanoma treatment, but here it is observed in primary cells. In comparison, cluster 7, which also showed features of UPR activation and MYC pathway induction, represented a more mixed and less coherent response. Cluster 7 contained a variety of compounds at high doses, including dabrafenib, but lacked the pathway specificity and transcriptional consistency observed in cluster 12.

By contrast, clusters 3, 5, and 9 encompassed compounds such as colchicine, nocodazole, podophyllotoxin, and cevipabulin fumarate that produce a distinct, stress-adapted transcriptional signature (Figure 6*a,b*). Like cluster 12, these clusters suppressed differentiation-related pathways (myogenesis, EMT, KRAS signaling), but were distinguished by strong upregulation of G2M checkpoint and DNA damage response signatures (Figure 6*b*). This pattern reflected a phenotype of mitotic stress and checkpoint activation, a cytostatic state in which melanocytes deprioritize biosynthesis and lineage-specific programs in favor of genomic integrity maintenance. These profiles may capture early transcriptional consequences of drugs such as colchicine, nocodazole, podophyllotoxin, and cevipabulin fumarate, which target tubulin or inhibit myotube formation, or are DNA-damaging agents, underscoring their relevance in modeling pre-apoptotic or senescent states.

The dose–response curves shown in Figure 6*c* further supported these findings, highlighting dose-dependent upregulation of *HSPA9*, a mitochondrial heat shock protein that plays a key role in the UPR and cellular adaptation to proteotoxic stress. Simultaneously, downregulation of *PRRX1* reinforces the suppression of KRAS-related signaling programs.

### 2.4 HDAC inhibitor transcriptomic signatures recapitulate external data

Quantitative analysis of dose-dependent genes across all compounds revealed that HDAC inhibitors consistently produced extensive transcriptional changes in all four cell types (Figure 7*a,b*). Known HDAC inhibitors TSA and panobinostat induced significant dose-dependent expression changes in over 3,000 genes per cell type, highlighting their potent epigenetic regulatory effects. Compounds with the most dose-dependent genes were TSA and another control compound, BFA. Other compound classes with substantial dose-dependent gene signatures included cytotoxic stressors (topotecan hydrochloride hydrate, (S)-(+)-camptothecin, idarubicin, mitoxantrone, cycloheximide, emetine dihydrochloride hydrate, and MPP⁺), stress pathway activators (thapsigargin, calcimycin, auranofin, and CCT018159), signal transduction modulators (torin2, LY2109761, CEP-33779, ouabain, hydrocortisone, and corticosterone), and cytoskeletal disruptors/mitosis inhibitors (rigosertib and cevipabulin fumarate).

**Figure 7:**
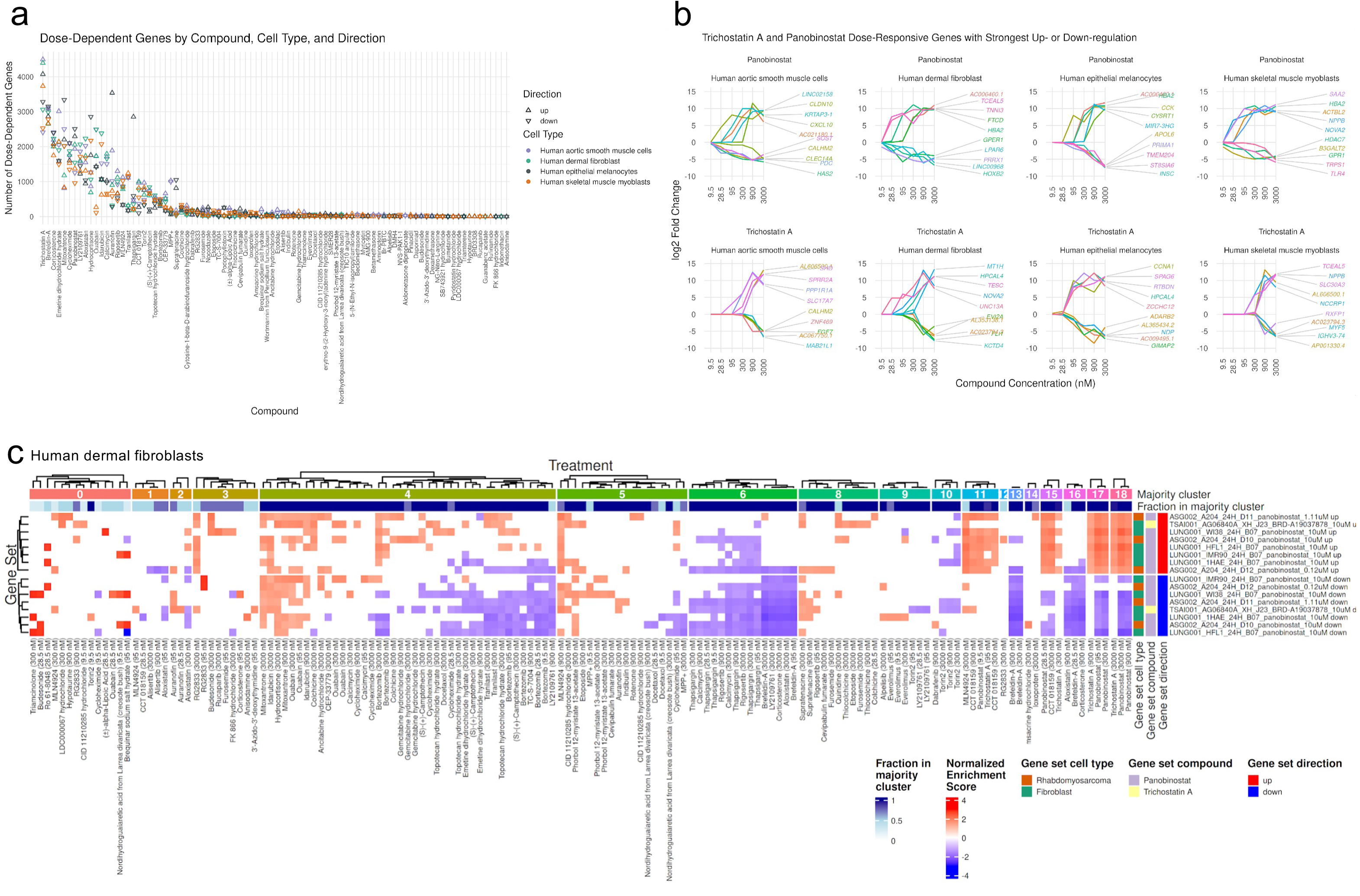
(*a*) Number of dose-dependent genes in our study by compound and cell type (*b*) Top 5 and bottom 5 dose-dependent gene-expression changes for trichostatin A and panobinostat across all cell types. (*c*) Gene set enrichment analysis for human dermal fibroblasts treated with trichostatin A and panobinostat using gene sets derived from the L1000 dataset.

In order to benchmark our findings, we compared our data against established transcriptomic drug perturbation signatures from the L1000 dataset. In a similar fashion to the MsigDB enrichment analysis, we performed enrichment analysis of our HDAC inhibitors (TSA and panobinostat) against gene sets derived from the L1000 dataset in related cell lineages treated with the same compounds (Duan et al., 2016; Subramanian et al., 2017). This analysis revealed significant concordance between our primary cell responses and those observed in the L1000 dataset across multiple cell types treated with TSA and panobinostat (Figure 7*c*, Supplementary Figures 3,4). Hierarchical clustering analysis of gene sets enriched for HDAC inhibitor response revealed that TSA and panobinostat clustered together in a concentration-dependent manner for all cell types. In the clustering pattern in dermal fibroblasts, we observed three distinct groups corresponding to low (clusters 11), middle (clusters 15 and 17), and high concentrations (cluster 18), demonstrating the dose-responsive nature of these compounds (Figure 7*c*). Additionally, CCT018159, which may also act as an HDAC inhibitor, was clustered with TSA and panobinostat in clusters 11 and 15 (Pinzi et al., 2020). This concentration-dependent clustering was also observed in the other three cell types (Supplementary Figures 3,4) and validates the sensitivity of our DRUG-seq approach in capturing graded transcriptional responses. The high degree of similarity between our DRUG-seq profiles and the L1000 signatures for these well-characterized compounds provided external validation of our experimental approach while highlighting the value of primary cells for capturing physiologically relevant drug responses.

## 3. Discussion and Future Directions

This study demonstrates the utility of high-throughput DRUG-seq in physiologically-relevant primary human cell models—AoSMCs, SkMMs, dermal fibroblasts, and melanocytes—to capture cell type-specific transcriptional responses to pharmacological perturbations. By profiling 89 compounds across six concentrations, we created a scalable and versatile resource that resolves transcriptional changes at both the gene and pathway levels. DRUG-seq effectively distinguishes cell types based on global gene expression profiles, captures dose-dependent effects, and identifies both conserved and distinct transcriptional signatures. These include canonical responses to known perturbations (e.g., BFA’s pan-cellular disruption of protein secretion, TSA’s dose-dependent chromatin modulation, and DEX’s selective hypoxia modulation in AoSMCs), as well as more nuanced, cell type-specific effects such as GR antagonist-induced anabolic signaling in fibroblasts and EMT pathway activation in SkMMs. Beyond validating known compound responses as we have shown here, DRUG-seq data can be further interrogated to classify compounds with previously unrecognized MOAs, such as CCT018159 possibly acting as an HDAC inhibitor. Moreover, this dataset is well-suited for queries against transcriptional disease signatures, facilitating the identification of compounds that reverse disease states or that can be repurposed for new therapeutic indications.

Other compound perturbation datasets with transcriptomic readout have been previously generated, such as the LINCS dataset using the L1000 platform, numerous single-cell datasets and, more recently, the Tahoe-100M dataset (Duan et al., 2016; Peidli et al., 2024; Zhang et al., 2025). While these datasets are invaluable for characterizing response to compounds, the DRUG-seq platform we have described here has several advantages. First, this framework is ideally suited for scaling to large chemical libraries of tens of thousands of compounds that can be screened individually or using combinatorial perturbations. Second, the choice of disease-relevant primary human cell types with diverse genetic backgrounds enables the generation of context-specific and clinically relevant datasets capturing regulatory nuances essential for understanding therapeutic responses (Schwanhäusser et al., 2011). Third, higher gene and transcript recovery compared to single-cell datasets provides enhanced characterization of the transcriptome. Fourth, this platform’s compatibility with multiwell formats allows for potential integration with other multiomic and functional assays, further expanding its utility.

Given these advantages, our DRUG-seq dataset represents a valuable resource for training and benchmarking emerging AI models designed to predict cellular mechanisms and perturbation–response relationships (Bunne et al., 2024; Lotfollahi et al., 2023; Qi et al., 2024; Roohani et al., 2024). These models aim to generalize drug effects across unseen cell types or conditions, which can be critical for applications in MOA inference, drug repurposing, and personalized medicine. However, many of these tools are currently trained on datasets with limited cell type diversity, low transcriptome coverage, or reliance on immortalized lines. In contrast, our DRUG-seq platform offers high-resolution, transcriptionally rich profiles from many different cell types, including primary human cells with diverse lineages and states, enabling more accurate modeling of cell state dynamics and pharmacological effects.

As more high-quality perturbation datasets are generated using platforms like DRUG-seq, the accuracy and biological fidelity of context-aware predictive models will correspondingly improve. Ultimately, this integration will expand the reach of *in silico* drug discovery, enable targeted hypothesis generation for disease-relevant pathways, and increase the clinical translatability of computational pharmacology.

## 4. Methods

### 4.1 Cell culture and treatment

Primary human aortic smooth muscle cells (C0075C), epithelial melanocytes (C0025C), SkMMs (A11440), and dermal fibroblasts (C0135C) were obtained from ThermoFisher Scientific. Cells were cultured in vendor-recommended media and maintained for no more than 10 passages before use. For screening, cells were seeded into 384-well plates using a BioTek 406FX liquid handler.

Compounds (Supplementary Table 1) were dispensed into wells using an Echo 525 acoustic dispenser in a randomized layout to minimize systematic bias. Each 384-well plate was incubated at 37°C with 5% CO_2_ in a Liconic static incubator for 24 hours. After incubation, cells were washed and lysed directly in the plates using a buffer containing 50 mM Tris-HCl pH 8.0, 75 mM KCl, 6% Ficoll PM-400, 0.15% Triton X-100, and 0.5 U/µL RNaseOUT. Lysed plates were stored at -80°C until library preparation.

A total of 89 pharmacologically-active compounds (some selected from the LOPAC® 1280 library, Sigma Aldrich, #12352202), were screened at six concentrations, with four replicates per condition. Three volumes of compounds (250 nL, 75 nL, and 25 nL) were dispensed into wells containing 40 µL of media, resulting in final DMSO concentrations of 0.625%, 0.1875%, and 0.0625%, respectively. DMSO controls were included on every plate in six replicates at each concentration. For each cell type, eight 384-well plates were screened for a total of 32 plates.

### 4.2 cDNA Library preparation and sequencing

RNA present in the lysate was converted to complementary DNA (cDNA) through reverse transcription (RT) as described in (Ye et al., 2018). The RT product was subsequently amplified (using instrument Mastercycler X50i), and full-length cDNA was quantified. The resulting cDNA was then used as input for Illumina DNA library preparation, employing custom primers designed to amplify the 3’ end of transcripts while preserving well barcodes and unique molecular identifiers (UMIs). Final libraries underwent quality control assessment via qPCR and Bioanalyzer before being sequenced on a NovaSeq X platform.

### 4.3 Read alignment and UMI counting

Following sequencing, raw reads were demultiplexed, and quality control checks were performed using FastQC (Andrews, 2012). Ribosomal RNA (rRNA) contamination was removed using the BBTools package bbduk to improve data specificity (Bushnell et al., 2017). Reads were then aligned to the hg38 reference genome using the STAR aligner, and PCR duplicates were removed using umi-tools (Dobin et al., 2013; Smith et al., 2017; The Genome Sequencing Consortium, 2001). De-duplicated transcript counts were then aggregated at the gene level to generate a UMI-per-gene matrix for each sample using the subread package (Shi et al., 2019).

### 4.4 QC, Sample filtering, PCA and differential expression analysis

Quality control (QC) metrics—total sequencing reads, the percentage of reads retained after rRNA removal, the percent of features mapped, the percentage of reads mapped to mitochondrial genes, sequencing saturation (1-(total_umi_count/total_sequenced_reads)) and number of genes with UMI>=3—are shown in Supplementary Figure 1. All control samples were sequenced to a depth of at least 2 million reads and exhibited tight distribution across other QC metrics. Ultimately, all controls for all four primary human cell types had an average gene (including all noncoding lncRNAs, protein coding and non-coding RNAs) recovery of just over 10,000 transcripts with greater than three unique UMI counts (Supplementary Figure 1), ensuring robust transcriptional data quality across plates.

LogCPM expression values were calculated using the Voom package with each condition consisting of a specific cell type, compound, and concentration (Law et al., 2014). To identify outliers, we first computed all pair-wise Euclidean distances between samples from the same condition using Voom normalized log2(CPM+0.5) values. We computed the mean distance of each sample from other samples in the same condition. Finally, samples with a mean distance greater than 1.5 times the inter-quartile range were flagged as outliers and excluded from downstream analyses.

For plate-to-plate correlation values, the mean genome-wide Pearson correlation coefficient was calculated for Voom normalized expression values between plates for the DMSO control wells (all concentrations) and for the positive control wells (TSA, BFA, and DEX), including all concentrations (Figure 2*a*).

Differential gene expression analysis was performed using the DESeq2 package, comparing samples treated with different concentrations of DMSO-solubilized compounds to samples treated with the same DMSO concentration (Love et al., 2014). Genes with adjusted p-value lower than 0.05 and absolute shrunken log2FoldChange value (calculated using the default apeglm method) greater than or equal to 1 were considered as “significantly differentially expressed” in downstream analyses. See Supplementary Data for DESeq2 results. PCA was performed on the top 500 most variable genes using Voom normalized expression values on all samples (Supplementary Figure 2).

### 4.5 Cell marker gene expression comparison

Cell marker genes were selected for each of the primary human cell types using two databases: the Chan-Zuckerberg Institute and the CellMarker 2.0 database (Abdulla et al., 2023; Hu et al., 2023). Using the internal search feature of each database, top cell marker genes were pulled out for “smooth muscle cells”, “skeletal muscle myoblasts”, “skin fibroblasts”, and “melanocytes”. The generated gene lists for all cell types from both databases were combined. Duplicate genes were removed and the gene list was further collapsed by selecting one or two representative genes from groups of gene families.

To compare expression across the four cell types, mean Voom normalized expression data for all wells containing the lowest DMSO concentration (0.0625%) for each cell type were visualized on a UMAP plot (Figure 2*b*). Cell type marker genes described above were highlighted on internal cell type data, using the R package pheatmap (Figure 2*c*) (Kolde, 2018).

### 4.6 Graph-based clustering and cluster annotation

Graph-based clustering and cluster visualization with UMAP were performed using Seurat v5 on the same set of samples used for differential expression analysis (Hafemeister & Satija, 2019; Hao et al., 2024). Specifically, the SCTransform method was used to normalize UMI counts and regress out DMSO concentration, and Canonical Correlation Analysis (CCA) was used to integrate data from different plates and correct for plate batch effects. Clustering was then performed using the default Seurat method, applying the Louvain algorithm (using a resolution value of 0.6) to a shared nearest neighbor graph computed from the top 30 CCA components. Finally, UMAP plots were generated using the same CCA components with Seurat default settings. The relatively low clustering resolution value was chosen to capture major trends, including similarities between compounds, although the dataset is complex enough to support much higher cluster resolutions if needed.

To identify expression patterns associated with each treatment, we performed gene set enrichment analysis using the ClusterProfiler GSEA function (with a p-value cutoff of 0.05 and minimum gene set size of 10) with shrunken log2 fold-change values obtained from DESeq2 differential expression analysis, and either MSigDB Hallmark gene sets or the l1000_cp gene sets derived from the L1000 transcriptomics dataset (Evangelista et al., 2022; Liberzon et al., 2015; Yu et al., 2012). In the latter case, we only considered gene sets for fibroblast, rhabdomyosarcoma, and melanoma cell lines, as proxies for primary fibroblasts, muscle cells, and melanocytes, respectively, treated with TSA (BRD-A19037878 or BRD-K68202742 compound IDs) or panobinostat. To improve visualization, we further subsetted L1000 melanoma gene sets by keeping only gene sets for 24h treatment, and only one randomly-selected gene set for each combination of cell type, compound, concentration, and timepoint. We then inspected enriched gene sets associated with each treatment and cluster by plotting heatmaps of normalized enrichment scores (NES) using the ComplexHeatmap package (Gu, 2022). Since fibroblasts, AoSMCs, and SkMMs share the same mesodermal origin, we reported, for each derived primary cell line, the enrichment for gene sets obtained from both fibroblast and rhabdomyosarcoma cell lines, also of mesodermal origin. Likewise, and for simplicity, primary melanocytes were compared to only melanoma, both being of ectodermal origin. The main cluster associated with each treatment was identified as the cluster containing the majority of samples receiving that treatment. Both the majority cluster and the fraction of samples mapping to it are reported on the heatmaps. See Supplementary Data for compound-cluster relationships and for GSEA results.

For each positive or negative gene set enrichment, we identified top contributing genes by aggregating the ranks of GSEA leading-edge genes across all treatments using the robust rank aggregation method and selecting the top 4 genes with the highest rank (Kolde et al., 2012).

### 4.7 Identification of genes with compound dose-dependent expression

Genes with compound dose-dependent expression were identified by fitting a 4-parameter logistic (i.e., symmetric sigmoid) function to log2fc ∼ log2(concentration) and applying the following criteria using the drda R package: 1) a non-NA log2FoldChange is computed by DESeq2 at all compound concentrations; 2) at least one of DESeq2-computed log2FoldChange values is greater than 1 for genes with positive slope or lower than -1 for genes with negative slope (with the slope sign derived from the fitted delta parameter of the logistic function); 3) the 4-parameter logistic model provides a better fit than a horizontal line according to the Akaike Information Criterion; 4) the p-value from a Likelihood Ratio Test comparing the 4-parameter logistic model to a horizontal line, adjusted for multiple testing by cell line and compound using the Bonferroni-Hochberg procedure, is less than 0.05 (Malyutina et al., 2023). See Supplementary Data for a full list of dose-dependent genes.

## Supporting information

CompoundLibrary

## 5. Data Availability

All Supplementary Data is available at https://datapoints.ginkgo.bio/functional-genomics/gdpx2-processed-data.

---

**Supplementary Figure 1:**
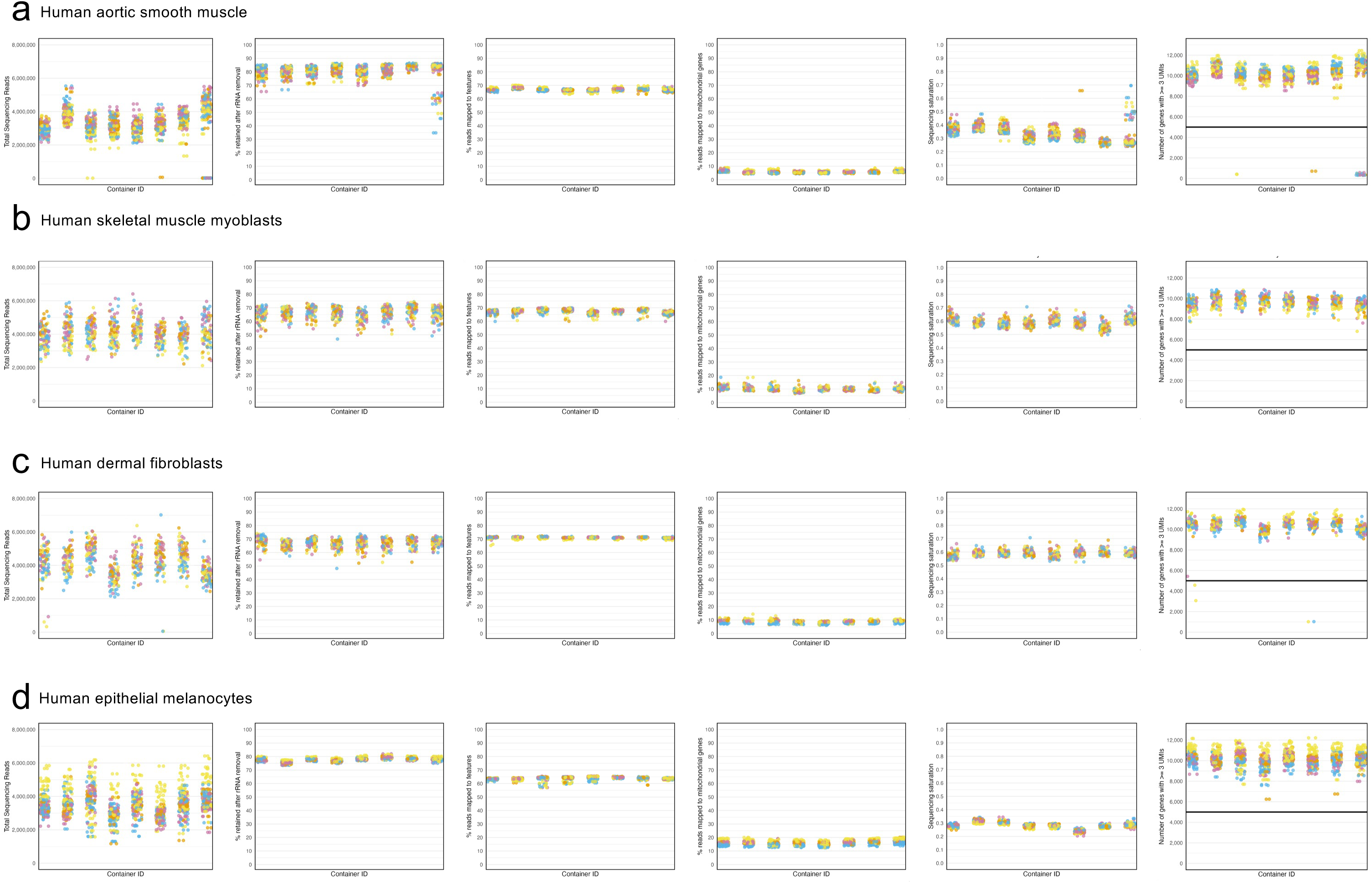
QC metrics showing, from left to right, total sequencing reads, the percentage of reads retained after rRNA removal, the percent of features mapped, the percentage of reads mapped to mitochondrial genes, sequencing saturation (1-(total_umi_count/total_sequenced_reads)) and number of genes with UMI>=3 across the 8, 384-well plates for (*a*) aortic smooth muscle cells (*b*) skeletal muscle myoblasts, *(c*) dermal fibroblasts, and (*d*) melanocytes.

**Supplementary Figure 2:**
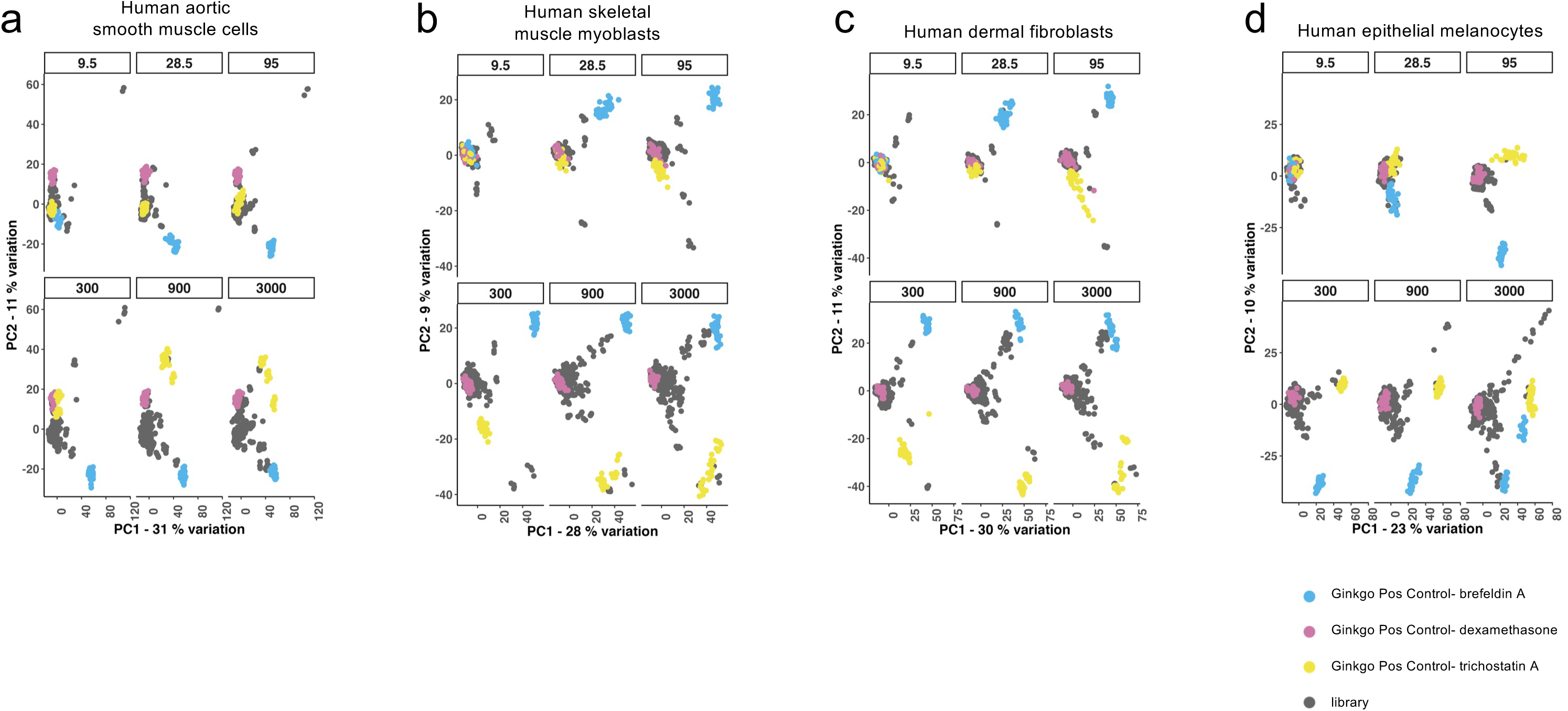
PCA by cell types separated by compound concentration. PCA analysis was run separately for each of the 6 compound concentrations (9.5, 28.5, 95, 300, 900, 3000 nM) and colored by positive control vs library. Generally, as compound concentration increased the control wells separated from the main cluster for each of the four cell types tested: (*a*) aortic smooth muscle cells, (*b*) skeletal muscle myoblasts, (*c*) dermal fibroblasts, and (*d*) melanocytes.

**Supplementary Figure 3:**
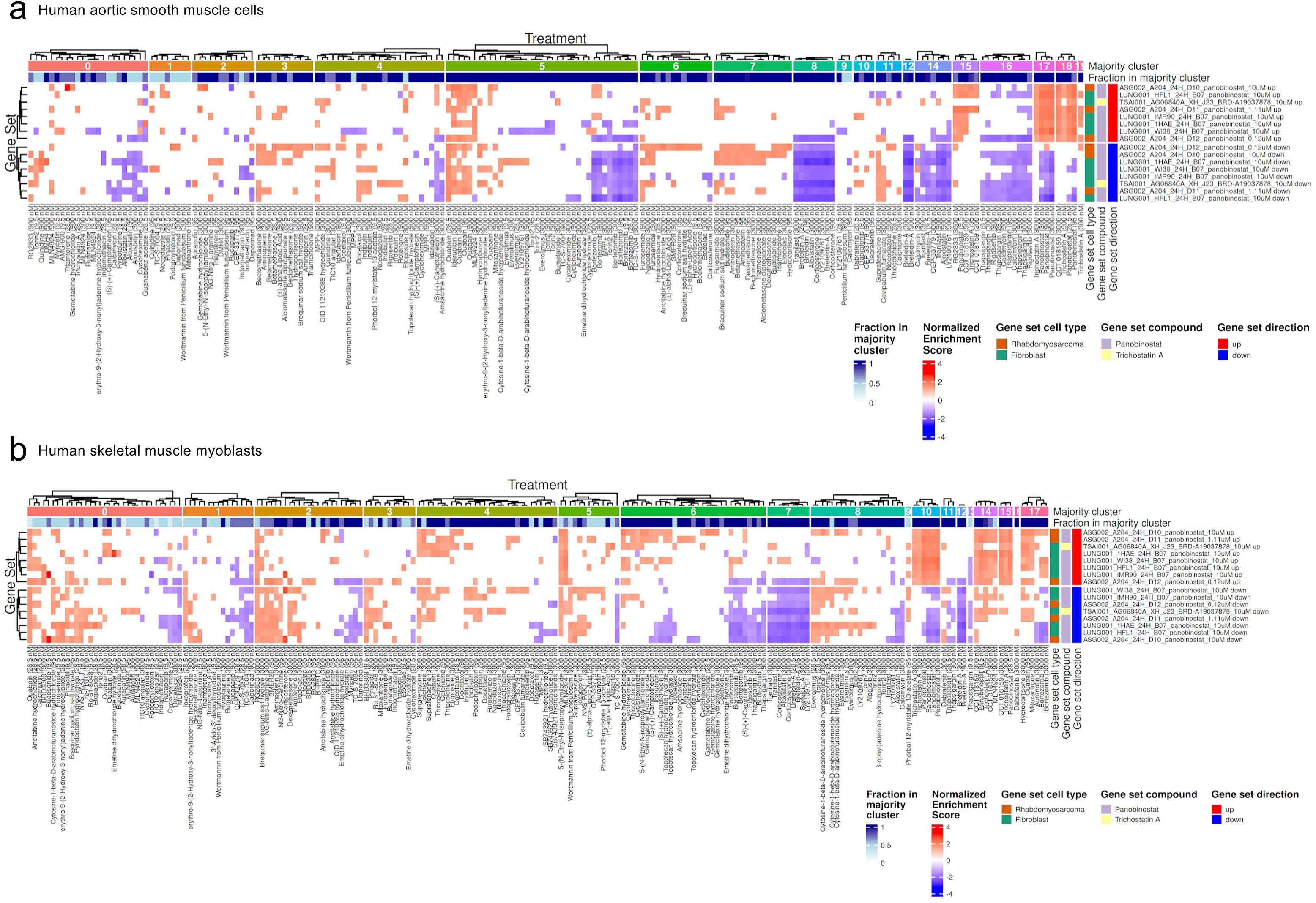
Gene set enrichment for (*a*) aortic smooth muscle cells and (*b*) skeletal muscle myoblasts against trichostatin A and panobinostat treated cells in L1000 dataset.

**Supplementary Figure 4:**
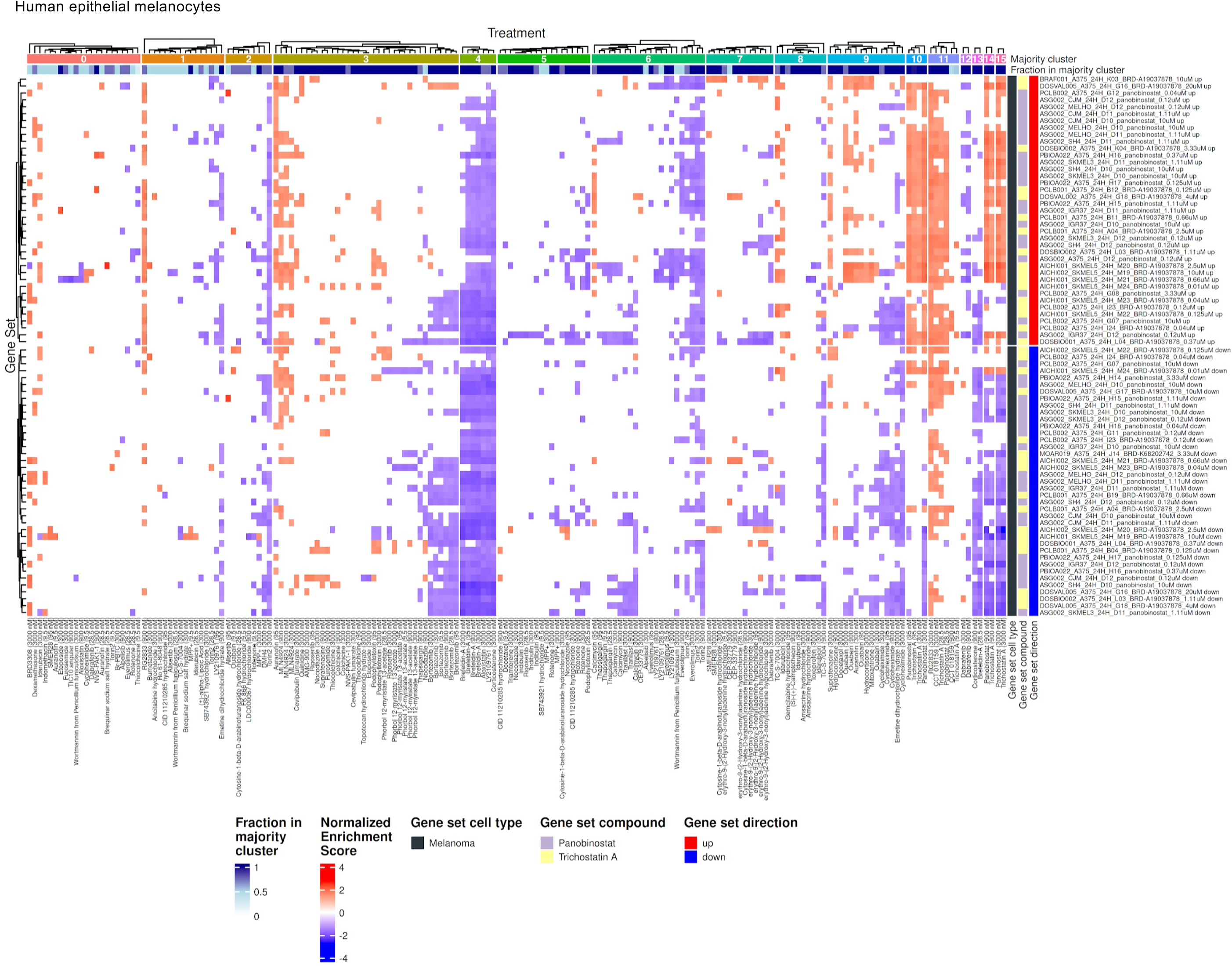
Gene set enrichment for melanocytes against trichostatin A and panobinostat treated cells in L1000 dataset.

## References

Abdulla, S., Aevermann, B., Assis, P., Badajoz, S., Bell, S. M., Bezzi, E., Cakir, B., Chaffer, J., Chambers, S., Cherry, J. M., Chi, T., Chien, J., Dorman, L., Garcia-Nieto, P., Gloria, N., Hastie, M., Hegeman, D., Hilton, J., Huang, T., … Carr, A. (2023). CZ CELL×GENE Discover: A single-cell data platform for scalable exploration, analysis and modeling of aggregated data (p. 2023.10.30.563174). bioRxiv. 10.1101/2023.10.30.563174

Andrews, S. (2012). Babraham Bioinformatics—FastQC A Quality Control tool for High Throughput Sequence Data. https://www.bioinformatics.babraham.ac.uk/projects/fastqc/

Bowyer, S., Lee, R., Fusi, A., & Lorigan, P. (2015). Dabrafenib and its use in the treatment of metastatic melanoma. Melanoma Management, 2(3), 199–208. 10.2217/mmt.15.21

Bundy, J. L., Everett, L. J., Rogers, J. D., Nyffeler, J., Byrd, G., Culbreth, M., Haggard, D. E., Word, L. J., Chambers, B. A., Davidson-Fritz, S., Harris, F., Willis, C., Paul-Friedman, K., Shah, I., Judson, R., & Harrill, J. A. (2024). High-Throughput Transcriptomics Screen of ToxCast Chemicals in U-2 OS Cells. Toxicology and Applied Pharmacology, 491, 117073. 10.1016/j.taap.2024.117073

Bunne, C., Roohani, Y., Rosen, Y., Gupta, A., Zhang, X., Roed, M., Alexandrov, T., AlQuraishi, M., Brennan, P., Burkhardt, D. B., Califano, A., Cool, J., Dernburg, A. F., Ewing, K., Fox, E. B., Haury, M., Herr, A. E., Horvitz, E., Hsu, P. D., … Quake, S. R. (2024). How to build the virtual cell with artificial intelligence: Priorities and opportunities. Cell, 187(25), 7045–7063. 10.1016/j.cell.2024.11.015

Bush, E. C., Ray, F., Alvarez, M. J., Realubit, R., Li, H., Karan, C., Califano, A., & Sims, P. A. (2017). PLATE-Seq for genome-wide regulatory network analysis of high-throughput screens. Nature Communications, 8(1), 105. 10.1038/s41467-017-00136-z

Bushnell, B., Rood, J., & Singer, E. (2017). BBMerge—Accurate paired shotgun read merging via overlap. PloS One, 12(10), e0185056. 10.1371/journal.pone.0185056

Butz, H., Saskői, É., Krokker, L., Vereczki, V., Alpár, A., Likó, I., Tóth, E., Szőcs, E., Cserepes, M., Nagy, K., Kacskovics, I., & Patócs, A. (2023). Context-Dependent Role of Glucocorticoid Receptor Alpha and Beta in Breast Cancer Cell Behaviour. Cells, 12(5), 784. 10.3390/cells12050784

Cho, D. S., & Doles, J. D. (2017). Single cell transcriptome analysis of muscle satellite cells reveals widespread transcriptional heterogeneity. Gene, 636, 54–63. 10.1016/j.gene.2017.09.014

Czuwara-Ladykowska, J., Makiela, B., Smith, E. A., Trojanowska, M., & Rudnicka, L. (2001). The inhibitory effects of camptothecin, a topoisomerase I inhibitor, on collagen synthesis in fibroblasts from patients with systemic sclerosis. Arthritis Research, 3(5), 311–318. 10.1186/ar321

Dai, W., Qiao, X., Fang, Y., Guo, R., Bai, P., Liu, S., Li, T., Jiang, Y., Wei, S., Na, Z., Xiao, X., & Li, D. (2024). Epigenetics-targeted drugs: Current paradigms and future challenges. Signal Transduction and Targeted Therapy, 9(1), 332. 10.1038/s41392-024-02039-0

Dobin, A., Davis, C. A., Schlesinger, F., Drenkow, J., Zaleski, C., Jha, S., Batut, P., Chaisson, M., & Gingeras, T. R. (2013). STAR: Ultrafast universal RNA-seq aligner. *Bioinformatics (Oxford*, England), 29(1), 15–21. 10.1093/bioinformatics/bts635

Dongre, A., & Weinberg, R. A. (2019). New insights into the mechanisms of epithelial-mesenchymal transition and implications for cancer. Nature Reviews. Molecular Cell Biology, 20(2), 69–84. 10.1038/s41580-018-0080-4

Duan, Q., Reid, S. P., Clark, N. R., Wang, Z., Fernandez, N. F., Rouillard, A. D., Readhead, B., Tritsch, S. R., Hodos, R., Hafner, M., Niepel, M., Sorger, P. K., Dudley, J. T., Bavari, S., Panchal, R. G., & Ma’ayan, A. (2016). L1000CDS2: LINCS L1000 characteristic direction signatures search engine. NPJ Systems Biology and Applications, 2, 16015-. 10.1038/npjsba.2016.15

Eastman, A. (2016). Improving anticancer drug development begins with cell culture: Misinformation perpetrated by the misuse of cytotoxicity assays. Oncotarget, 8(5), 8854–8866. 10.18632/oncotarget.12673

Evangelista, J. E., Clarke, D. J. B., Xie, Z., Lachmann, A., Jeon, M., Chen, K., Jagodnik, K. M., Jenkins, S. L., Kuleshov, M. V., Wojciechowicz, M. L., Schürer, S. C., Medvedovic, M., & Ma’ayan, A. (2022). SigCom LINCS: Data and metadata search engine for a million gene expression signatures. Nucleic Acids Research, 50(W1), W697–W709. 10.1093/nar/gkac328

Feeney, G. P., Errington, R. J., Wiltshire, M., Marquez, N., Chappell, S. C., & Smith, P. J. (2003). Tracking the cell cycle origins for escape from topotecan action by breast cancer cells. British Journal of Cancer, 88(8), 1310–1317. 10.1038/sj.bjc.6600889

Fujiwara, T., Oda, K., Yokota, S., Takatsuki, A., & Ikehara, Y. (1988). Brefeldin A causes disassembly of the Golgi complex and accumulation of secretory proteins in the endoplasmic reticulum. The Journal of Biological Chemistry, 263(34), 18545–18552.

Gu, Z. (2022). Complex heatmap visualization. iMeta, 1(3), e43. 10.1002/imt2.43

Hafemeister, C., & Satija, R. (2019). Normalization and variance stabilization of single-cell RNA-seq data using regularized negative binomial regression. Genome Biology, 20(1), 296. 10.1186/s13059-019-1874-1

Hao, Y., Stuart, T., Kowalski, M. H., Choudhary, S., Hoffman, P., Hartman, A., Srivastava, A., Molla, G., Madad, S., Fernandez-Granda, C., & Satija, R. (2024). Dictionary learning for integrative, multimodal and scalable single-cell analysis. Nature Biotechnology, 42(2), 293–304. 10.1038/s41587-023-01767-y

Hu, C., Li, T., Xu, Y., Zhang, X., Li, F., Bai, J., Chen, J., Jiang, W., Yang, K., Ou, Q., Li, X., Wang, P., & Zhang, Y. (2023). CellMarker 2.0: An updated database of manually curated cell markers in human/mouse and web tools based on scRNA-seq data. Nucleic Acids Research, 51(D1), D870–D876. 10.1093/nar/gkac947

Kawami, M., Harada, R., Ojima, T., Yamagami, Y., Yumoto, R., & Takano, M. (2019). Association of cell cycle arrest with anticancer drug-induced epithelial-mesenchymal transition in alveolar epithelial cells. Toxicology, 424, 152231. 10.1016/j.tox.2019.06.002

King, A. J., Arnone, M. R., Bleam, M. R., Moss, K. G., Yang, J., Fedorowicz, K. E., Smitheman, K. N., Erhardt, J. A., Hughes-Earle, A., Kane-Carson, L. S., Sinnamon, R. H., Qi, H., Rheault, T. R., Uehling, D. E., & Laquerre, S. G. (2013). Dabrafenib; Preclinical Characterization, Increased Efficacy when Combined with Trametinib, while BRAF/MEK Tool Combination Reduced Skin Lesions. PLoS ONE, 8(7), e67583. 10.1371/journal.pone.0067583

Kolde, R. (2018). pheatmap: Pretty Heatmaps. https://raivokolde.r-universe.dev/pheatmap

Kolde, R., Laur, S., Adler, P., & Vilo, J. (2012). Robust rank aggregation for gene list integration and meta-analysis. Bioinformatics (Oxford, England), 28(4), 573–580. 10.1093/bioinformatics/btr709

La, J., Reed, E., Chan, L., Smolyaninova, L. V., Akomova, O. A., Mutlu, G. M., Orlov, S. N., & Dulin, N. O. (2016). Downregulation of TGF-β Receptor-2 Expression and Signaling through Inhibition of Na/K-ATPase. PLoS ONE, 11(12), e0168363. 10.1371/journal.pone.0168363

Lamb, J., Crawford, E. D., Peck, D., Modell, J. W., Blat, I. C., Wrobel, M. J., Lerner, J., Brunet, J.-P., Subramanian, A., Ross, K. N., Reich, M., Hieronymus, H., Wei, G., Armstrong, S. A., Haggarty, S. J., Clemons, P. A., Wei, R., Carr, S. A., Lander, E. S., & Golub, T. R. (2006). The Connectivity Map: Using gene-expression signatures to connect small molecules, genes, and disease. Science (New York, N.Y.), 313(5795), 1929–1935. 10.1126/science.1132939

Lamouille, S., Xu, J., & Derynck, R. (2014). Molecular mechanisms of epithelial-mesenchymal transition. Nature Reviews. Molecular Cell Biology, 15(3), 178–196. 10.1038/nrm3758

Law, C. W., Chen, Y., Shi, W., & Smyth, G. K. (2014). voom: Precision weights unlock linear model analysis tools for RNA-seq read counts. Genome Biology, 15(2), R29. 10.1186/gb-2014-15-2-r29

Li, H., Qiu, J., & Fu, X.-D. (2012). RASL-seq for Massively Parallel and Quantitative Analysis of Gene Expression. Current Protocols in Molecular Biology, 98(1), 4.13.1–4.13.9. 10.1002/0471142727.mb0413s98

Liberzon, A., Birger, C., Thorvaldsdóttir, H., Ghandi, M., Mesirov, J. P., & Tamayo, P. (2015). The Molecular Signatures Database Hallmark Gene Set Collection. Cell Systems, 1(6), 417–425. 10.1016/j.cels.2015.12.004

Lotfollahi, M., Klimovskaia Susmelj, A., De Donno, C., Hetzel, L., Ji, Y., Ibarra, I. L., Srivatsan, S. R., Naghipourfar, M., Daza, R. M., Martin, B., Shendure, J., McFaline-Figueroa, J. L., Boyeau, P., Wolf, F. A., Yakubova, N., Günnemann, S., Trapnell, C., Lopez-Paz, D., & Theis, F. J. (2023). Predicting cellular responses to complex perturbations in high-throughput screens. Molecular Systems Biology, 19(6), e11517. 10.15252/msb.202211517

Love, M. I., Huber, W., & Anders, S. (2014). Moderated estimation of fold change and dispersion for RNA-seq data with DESeq2. Genome Biology, 15(12), 550. 10.1186/s13059-014-0550-8

Malyutina, A., Tang, J., & Pessia, A. (2023). drda: An R Package for Dose-Response Data Analysis Using Logistic Functions. Journal of Statistical Software, 106, 1–26. 10.18637/jss.v106.i04

Oakley, R. H., & Cidlowski, J. A. (2013). The biology of the glucocorticoid receptor: New signaling mechanisms in health and disease. The Journal of Allergy and Clinical Immunology, 132(5), 1033–1044. 10.1016/j.jaci.2013.09.007

Peidli, S., Green, T. D., Shen, C., Gross, T., Min, J., Garda, S., Yuan, B., Schumacher, L. J., Taylor-King, J. P., Marks, D. S., Luna, A., Blüthgen, N., & Sander, C. (2024). scPerturb: Harmonized single-cell perturbation data. Nature Methods, 21(3), 531–540. 10.1038/s41592-023-02144-y

Pinzi, L., Benedetti, R., Altucci, L., & Rastelli, G. (2020). Design of Dual Inhibitors of Histone Deacetylase 6 and Heat Shock Protein 90. ACS Omega, 5(20), 11473–11480. 10.1021/acsomega.0c00559

Qi, X., Zhao, L., Tian, C., Li, Y., Chen, Z.-L., Huo, P., Chen, R., Liu, X., Wan, B., Yang, S., & Zhao, Y. (2024). Predicting transcriptional responses to novel chemical perturbations using deep generative model for drug discovery. Nature Communications, 15(1), 9256. 10.1038/s41467-024-53457-1

Reil, T. D., Sarkar, R., Kashyap, V. S., Sarkar, M., & Gelabert, H. A. (1999). Dexamethasone suppresses vascular smooth muscle cell proliferation. The Journal of Surgical Research, 85(1), 109–114. 10.1006/jsre.1999.5665

Rhen, T., & Cidlowski, J. A. (2005). Antiinflammatory Action of Glucocorticoids—New Mechanisms for Old Drugs. New England Journal of Medicine, 353(16), 1711–1723. 10.1056/NEJMra050541

Roohani, Y., Huang, K., & Leskovec, J. (2024). Predicting transcriptional outcomes of novel multigene perturbations with GEARS. Nature Biotechnology, 42(6), 927–935. 10.1038/s41587-023-01905-6

Sandbo, N., Ngam, C., Torr, E., Kregel, S., Kach, J., & Dulin, N. (2013). Control of Myofibroblast Differentiation by Microtubule Dynamics through a Regulated Localization of mDia2. The Journal of Biological Chemistry, 288(22), 15466–15473. 10.1074/jbc.M113.464461

Schwanhäusser, B., Busse, D., Li, N., Dittmar, G., Schuchhardt, J., Wolf, J., Chen, W., & Selbach, M. (2011). Global quantification of mammalian gene expression control. Nature, 473(7347), 337–342. 10.1038/nature10098

Shi, W., Liao, Y., & Smyth, G. (2019). Rsubread. Bioconductor. http://bioconductor.org/packages/Rsubread/

Shin, K. W. D., Atalay, M. V., Cetin-Atalay, R., O’Leary, E. M., Glass, M. E., Szafran, J. C. H., Woods, P. S., Meliton, A. Y., Shamaa, O. R., Tian, Y., Mutlu, G. M., & Hamanaka, R. B. (2025). mTOR signaling regulates multiple metabolic pathways in human lung fibroblasts after TGF-β and in pulmonary fibrosis. American Journal of Physiology. Lung Cellular and Molecular Physiology, 328(2), L215–L228. 10.1152/ajplung.00189.2024

Smith, T., Heger, A., & Sudbery, I. (2017). UMI-tools: Modeling sequencing errors in Unique Molecular Identifiers to improve quantification accuracy. Genome Research, 27(3), 491–499. 10.1101/gr.209601.116

Subramanian, A., Narayan, R., Corsello, S. M., Peck, D. D., Natoli, T. E., Lu, X., Gould, J., Davis, J. F., Tubelli, A. A., Asiedu, J. K., Lahr, D. L., Hirschman, J. E., Liu, Z., Donahue, M., Julian, B., Khan, M., Wadden, D., Smith, I. C., Lam, D., … Golub, T. R. (2017). A Next Generation Connectivity Map: L1000 Platform and the First 1,000,000 Profiles. Cell, 171(6), 1437–1452.e17. 10.1016/j.cell.2017.10.049

The Genome Sequencing Consortium. (2001, February 15). Initial sequencing and analysis of the human genome. Nature. https://www.ncbi.nlm.nih.gov/assembly/88331

Voloshin, N., Tyurin-Kuzmin, P., Karagyaur, M., Akopyan, Z., & Kulebyakin, K. (2023). Practical Use of Immortalized Cells in Medicine: Current Advances and Future Perspectives. International Journal of Molecular Sciences, 24(16), 12716. 10.3390/ijms241612716

Yang, C., Tian, C., Hoffman, T. E., Jacobsen, N. K., & Spencer, S. L. (2021). Melanoma subpopulations that rapidly escape MAPK pathway inhibition incur DNA damage and rely on stress signalling. Nature Communications, 12(1), 1747. 10.1038/s41467-021-21549-x

Ye, C., Ho, D. J., Neri, M., Yang, C., Kulkarni, T., Randhawa, R., Henault, M., Mostacci, N., Farmer, P., Renner, S., Ihry, R., Mansur, L., Keller, C. G., McAllister, G., Hild, M., Jenkins, J., & Kaykas, A. (2018). DRUG-seq for miniaturized high-throughput transcriptome profiling in drug discovery. Nature Communications, 9(1), 4307. 10.1038/s41467-018-06500-x

Yoshida, M., Kijima, M., Akita, M., & Beppu, T. (1990). Potent and specific inhibition of mammalian histone deacetylase both in vivo and in vitro by trichostatin A. The Journal of Biological Chemistry, 265(28), 17174–17179.

Yu, G., Wang, L.-G., Han, Y., & He, Q.-Y. (2012). clusterProfiler: An R Package for Comparing Biological Themes Among Gene Clusters. OMICS : A Journal of Integrative Biology, 16(5), 284–287. 10.1089/omi.2011.0118

Zhang, J., Ubas, A. A., Borja, R. de, Svensson, V., Thomas, N., Thakar, N., Lai, I., Winters, A., Khan, U., Jones, M. G., Tran, V., Pangallo, J., Papalexi, E., Sapre, A., Nguyen, H., Sanderson, O., Nigos, M., Kaplan, O., Schroeder, S., … Yu, J. (2025). Tahoe-100M: A Giga-Scale Single-Cell Perturbation Atlas for Context-Dependent Gene Function and Cellular Modeling (p. 2025.02.20.639398). bioRxiv. 10.1101/2025.02.20.639398 (8 compounds on last plate)

