## Supplementary material for "Mapping the Transcriptional Landscape of Drug Responses in Primary Human Cells Using High-Throughput DRUG-seq": CompoundLibrary

| Drug | Type | Category | Function |
|---|---|---|---|
| Brefeldin-A | positive control | ER Stress and Calcium Homeostasis | Inhibitor of protein transport from the endoplasmic reticulum |
| Dexamethasone | positive control | Anti-inflammatory, Analgesic, and Fibrosis Modulation Agents | Corticosteroid |
| Trichostatin A | positive control | Antineoplastic Agents | HDAC inhibitor |
| DMSO | negative control | Miscellaneous | Solvent with anti-inflammatory and antioxidant properties |
| (S)-(+)-Camptothecin | library | Antineoplastic Agents | Topoisomerase I inhibitor |
| 3'-Azido-3'-deoxythymidine | library | DNA Repair and Metabolic Modulation | Antiviral agent, affects DNA synthesis |
| 5-(N-Ethyl-N-isopropyl)amiloride | library | Ion Channel Modulators | Sodium-hydrogen exchange inhibitor |
| Alclometasone dipropionate | library | Anti-inflammatory, Analgesic, and Fibrosis Modulation Agents | Corticosteroid |
| Alisertib | library | Antineoplastic Agents | Aurora kinase A inhibitor |
| Aloxistatin | library | Protein Synthesis and Degradation Inhibitors | Cysteine protease inhibitor |
| Alpelisib | library | Kinase Inhibitors | PI3K inhibitor |
| AMG-900 | library | Kinase Inhibitors | Aurora kinase inhibitor |
| Aminopterin | library | Antineoplastic Agents | Antifolate, inhibits dihydrofolate reductase |
| Amsacrine hydrochloride | library | Antineoplastic Agents | DNA intercalator, topoisomerase II inhibitor |
| Ancitabine hydrochloride | library | Antineoplastic Agents | Antineoplastic agent |
| Anisodamine | library | Anti-inflammatory, Analgesic, and Fibrosis Modulation Agents | Muscarinic acetylcholine receptor antagonist |
| Auranofin | library | Anti-inflammatory, Analgesic, and Fibrosis Modulation Agents | Anti-rheumatic agent |
| Beclomethasone | library | Anti-inflammatory, Analgesic, and Fibrosis Modulation Agents | Corticosteroid |
| Betamethasone | library | Anti-inflammatory, Analgesic, and Fibrosis Modulation Agents | Corticosteroid |
| Bortezomib | library | Protein Synthesis and Degradation Inhibitors | Proteasome inhibitor |
| Br-PBTC | library | Miscellaneous | Potential therapeutic agent |
| BRD3308 | library | Miscellaneous | Potential therapeutic agent |
| Brequinar sodium salt hydrate | library | DNA Repair and Metabolic Modulation | Inhibitor of dihydroorotate dehydrogenase, affecting pyrimidine synthesis |
| Budesonide | library | Anti-inflammatory, Analgesic, and Fibrosis Modulation Agents | Corticosteroid |
| Bumetanide | library | Ion Channel Modulators | Loop diuretic, inhibits Na-K-Cl cotransporter |
| Calcimycin | library | ER Stress and Calcium Homeostasis | Calcium ionophore |
| CCT 018159 | library | Kinase Inhibitors | CHK1 inhibitor |
| CEP-33779 | library | Kinase Inhibitors | JAK2 inhibitor |

| | | | |
|---|---|---|---|
| Cevipabulin fumarate | library | Antineoplastic Agents | Microtubule polymerization inhibitor |
| CID 11210285 hydrochloride | library | Miscellaneous | Potential therapeutic agent |
| Colchicine | library | Antineoplastic Agents | Microtubule polymerization inhibitor |
| Corticosterone | library | Miscellaneous | Glucocorticoid, involved in stress response |
| Cycloheximide | library | Protein Synthesis and Degradation Inhibitors | Protein synthesis inhibitor |
| Cytosine-1-beta-D-arabinofuranoside hydrochloride | library | Antineoplastic Agents | Antineoplastic agent, inhibits DNA synthesis |
| Dabrafenib | library | Kinase Inhibitors | BRAF inhibitor |
| Daporinad | library | DNA Repair and Metabolic Modulation | NAMPT inhibitor |
| DMH4 | library | Kinase Inhibitors | ALK5 inhibitor |
| Docetaxol | library | Antineoplastic Agents | Microtubule stabilizer |
| Emetine dihydrochloride hydrate | library | Protein Synthesis and Degradation Inhibitors | Protein synthesis inhibitor |
| erythro-9-(2-Hydroxy-3-nonyl)adenine hydrochloride | library | Miscellaneous | Adenosine deaminase inhibitor |
| Etodolac | library | Anti-inflammatory, Analgesic, and Fibrosis Modulation Agents | Nonsteroidal anti-inflammatory drug (NSAID) |
| Etoposide | library | Antineoplastic Agents | Topoisomerase II inhibitor |
| Everolimus | library | Kinase Inhibitors | mTOR inhibitor |
| Furosemide | library | Ion Channel Modulators | Loop diuretic, inhibits Na-K-Cl cotransporter |
| Gemcitabine hydrochloride | library | Antineoplastic Agents | Antineoplastic agent, inhibits DNA synthesis |
| Guanabenz acetate | library | Miscellaneous | Alpha-2 adrenergic agonist |
| Hydrocortisone | library | Anti-inflammatory, Analgesic, and Fibrosis Modulation Agents | Corticosteroid |
| Hypotaurine | library | DNA Repair and Metabolic Modulation | Antioxidant, sulfur-containing amino acid derivative |
| Idarubicin | library | Antineoplastic Agents | DNA intercalator, topoisomerase II inhibitor |
| Indibulin | library | Antineoplastic Agents | Microtubule polymerization inhibitor |
| Indomethacin | library | Anti-inflammatory, Analgesic, and Fibrosis Modulation Agents | Nonsteroidal anti-inflammatory drug (NSAID) |
| LDC000067 hydrochloride | library | Kinase Inhibitors | CDK9 inhibitor |
| Loxoprofen | library | Anti-inflammatory, Analgesic, and Fibrosis Modulation Agents | Nonsteroidal anti-inflammatory drug (NSAID) |
| LY2109761 | library | Kinase Inhibitors | TGF-beta receptor I/II kinase inhibitor |
| Mitoxantrone | library | Antineoplastic Agents | DNA intercalator, topoisomerase II inhibitor |
| MLN4924 | library | Antineoplastic Agents | NEDD8-activating enzyme inhibitor |
| MPP+ | library | DNA Repair and Metabolic Modulation | Neurotoxin, affects mitochondrial function |

| | | | |
|---|---|---|---|
| Myricetin | library | DNA Repair and Metabolic Modulation | Flavonoid with antioxidant properties |
| NG-Nitro-L-arginine | library | DNA Repair and Metabolic Modulation | Nitric oxide synthase inhibitor |
| Nocodazole | library | Antineoplastic Agents | Microtubule polymerization inhibitor |
| Nordihydroguaiaretic acid from Larrea divaricata (creosote bush) | library | DNA Repair and Metabolic Modulation | Antioxidant, lipoxygenase inhibitor |
| NVS-PAK1-1 | library | Kinase Inhibitors | PAK1 inhibitor |
| Ouabain | library | Ion Channel Modulators | Na+/K+-ATPase inhibitor |
| Panobinostat | library | Antineoplastic Agents | HDAC inhibitor |
| Phorbol 12-myristate 13-acetate | library | Kinase Inhibitors | Protein kinase C activator |
| Pinacidil | library | Ion Channel Modulators | Potassium channel opener |
| Podophyllotoxin | library | Antineoplastic Agents | Microtubule inhibitor |
| Pyridostatin hydrochloride | library | DNA Repair and Metabolic Modulation | G-quadruplex stabilizer |
| Quinidine | library | Ion Channel Modulators | Antiarrhythmic, Na+ channel blocker |
| RG2833 | library | Kinase Inhibitors | HDAC inhibitor |
| Rigosertib | library | Kinase Inhibitors | PLK1 inhibitor |
| Ro 61-8048 | library | DNA Repair and Metabolic Modulation | Kynurenine 3-monooxygenase inhibitor |
| Rotenone | library | DNA Repair and Metabolic Modulation | Mitochondrial complex I inhibitor |
| Rucaparib | library | Antineoplastic Agents | PARP inhibitor |
| Rufinamide | library | Anti-inflammatory, Analgesic, and Fibrosis Modulation Agents | Antiseizure medication with potential anti-inflammatory effects |
| SB743921 hydrochloride | library | Antineoplastic Agents | Kinesin spindle protein inhibitor |
| SMER28 | library | Protein Synthesis and Degradation Inhibitors | Autophagy inducer |
| Suprafenacine | library | Miscellaneous | Potential therapeutic agent |
| TC-S-7004 | library | Kinase Inhibitors | JAK2 inhibitor |
| Thapsigargin | library | ER Stress and Calcium Homeostasis | SERCA inhibitor |
| Thiocolchicine | library | Antineoplastic Agents | Microtubule polymerization inhibitor |
| TIC10 angular | library | Antineoplastic Agents | TRAIL-inducing compound |
| Topotecan hydrochloride hydrate | library | Antineoplastic Agents | Topoisomerase I inhibitor |
| Torin2 | library | Kinase Inhibitors | mTOR inhibitor |
| Tranilast | library | Anti-inflammatory, Analgesic, and Fibrosis Modulation Agents | Anti-fibrotic agent |
| Triamcinolone | library | Anti-inflammatory, Analgesic, and Fibrosis Modulation Agents | Corticosteroid |
| Triamterene | library | Ion Channel Modulators | Potassium-sparing diuretic, inhibits epithelial sodium channels |

| | | | |
|---|---|---|---|
| Wortmannin from Penicillium funiculosum | library | Kinase Inhibitors | PI3K inhibitor |
| (¬±)-alpha-Lipoic Acid | library | DNA Repair and Metabolic Modulation | Antioxidant, affects mitochondrial function |
| FK 866 hydrochloride | library | DNA Repair and Metabolic Modulation | NAMPT inhibitor |
